# Enhanced environmental complexity worsens experimental colitis and dysregulates microbiota-gut-brain axis signalling in female mice

**DOI:** 10.64898/2026.06.26.734703

**Authors:** Giulia Petracco, Isabella Faimann, Eva Gruden, Melanie Kienzl, Elmar Zügner, Fernanda Monedeiro, Christina Kumpitsch, Eva Tatzl, Günther Rauter, Sabine Obermüller, Thomas Altendorfer-Kroath, Christine Moissl-Eichinger, Rudolf Schicho, Christoph Magnes, Florian Reichmann

## Abstract

Ulcerative colitis (UC) is a chronic inflammatory disease characterized by colonic inflammation and bloody diarrhoea. Accumulating evidence suggests that UC not only affects the intestinal tract, but also distant organs including the brain. Environmental factors are key determinants of the disease course, yet the impact and potential disease modifying effects of living environment complexity on microbiota-gut-brain axis signalling during colitis remain unclear. To address this gap, we investigated how enhanced environmental complexity (EC) affects the disease course and gut-brain axis signalling during experimental colitis in mice. Our results show that EC exacerbates dextran sulphate sodium (DSS)-induced colitis in female mice, but not in male mice, as evidenced by greater weight loss and higher disease activity. Immune cell profiling across the gut-brain axis reveals strong effects of DSS treatment on colonic, circulating and brain immune cell populations and a restriction of central nervous system (CNS) T cell infiltration due to EC. In addition, female EC/DSS mice have higher circulating corticosterone levels than controls indicating chronic stress. Metabolomics across the gut-brain axis revealed that EC exacerbates colitis-induced metabolite perturbations in plasma, brain tissue, brain interstitial and cerebrospinal fluid. Notably, microbiota-derived metabolites, including deoxycholic acid and trimethylamine-N-oxide (TMAO), are increased in EC/DSS mice, concordant with EC-associated microbiome changes and anxiety-like behaviour. Overall, this study indicates that EC worsens experimental colitis in female mice and directs microbiota-gut-brain axis signalling during colitis towards a less favourable state. From a translational perspective, this study highlights the importance of environmental factors for a sex-specific disease course of UC and associated neurobehavioral comorbidities.

**Highlights:**

- Enhanced environmental complexity (EC) exacerbates experimental colitis
- Colitis and EC have compartment-specific effects on immune cells
- EC augments colitis-induced metabolic shifts in plasma, brain and CSF
- Microbiota-derived metabolites are important players for the effects of EC

## Introduction

Inflammatory bowel disease (IBD) denotes a family of chronic gastrointestinal disorders, including the two major entities Crohn’s disease (CD) and ulcerative colitis (UC). These diseases are characterized by symptoms such as abdominal pain, diarrhoea and bloody stools, and they affect between five and ten million people worldwide (Fakhoury *et al*. 2014; Wang *et al*. 2023). In many cases, IBD not only involves the gut, but also affects other organs such as joints, eyes, skin, liver or even the brain, in so-called extraintestinal manifestations (Veloso 2011; Morís 2014; Rogler *et al*. 2021). Regarding the latter, it has been found that IBD patients are at a higher risk for mood disorders such as anxiety, depression and bipolar disorder (Bernstein *et al*. 2019; Bisgaard *et al*. 2022; Massironi *et al*. 2025; Petracco *et al*. 2025), suggesting a link between peripheral pathophysiological processes and neuropsychiatric disorders.

Psychological stress has been identified as a key factor in triggering IBD exacerbations (Fairbrass *et al*. 2022; Ge *et al*. 2022). It is known that chronic psychosocial and physical stress have a marked impact on both brain (McEwen *et al*. 2015) and gut function (Elsenbruch and Enck 2017), and that these effects are interrelated. Animal studies confirm these relationships, as experimental gastrointestinal inflammation, an endogenous stressor itself, alters emotional-affective behaviour and stress reactivity (Painsipp *et al*. 2011; Hassan *et al*. 2014; Reichmann *et al*. 2015; Nyuyki *et al*. 2018) and experimental psychosocial stress is known to aggravate inflammation and trigger relapses in animal models of IBD (Reber 2012). Furthermore, dysbiosis plays an important role in the pathogenesis of IBD (Iliev *et al*. 2025). Microbial factors are contributing to IBD-associated neuropsychiatric comorbidities via the microbiota-gut-brain (MGB) axis, a term that denotes the multi-directional communication between its components (Skonieczna-żydecka *et al*. 2018). For example, acute administration of dextran sodium sulphate (DSS) induces colitis, dysbiosis as well as anxiety and recognition memory deficits in mice, which can be normalized by *Lactobacillus*-based probiotics, directly linking intestinal inflammation, microbial shifts, and CNS function (Emge *et al*. 2016). In addition, when colitis is induced at weaning, mice exhibit long-lasting cognitive and anxiety-like deficits with hippocampal neuroinflammation and dysbiosis persisting into adulthood, demonstrating long-term MGB axis disruptions by early-life intestinal inflammation (Salvo *et al*. 2020). Targeting the MGB axis therapeutically has been suggested as a promising way to alleviate comorbid neurobehavioral disturbances and multiple potential therapeutics are under active investigation, such as faecal matter transplantation, psychobiotics, microbial metabolite supplementation and vagus nerve stimulation (Chen *et al*. 2018; Sinniger *et al*. 2020; Feng *et al*. 2023; Brown *et al*. 2024; Petracco *et al*. 2025)

While it is clear that adverse environmental factors, such as chronic stress, can markedly disrupt MGB axis signalling during IBD (Bisgaard *et al*. 2022; Bonaz *et al*. 2024), it remains to be determined whether putative beneficial environmental factors such as environmental enrichment exert an equally strong effect. Usually, enrichment refers to a number of procedures used in animal husbandry and scientific research aiming at the improvement of animal welfare and the promotion of species-specific behaviour under captivity (Kulpa-Eddy *et al*. 2005). These procedures typically involve more complex home cage environments (high environmental complexity, EC) with the aim to provide sufficient space for the expression of a wide spectrum of normal behaviours and the provision of natural and/or artificial objects to interact with, in order to stimulate mental and/or physical activity (Taylor *et al*. 2023). EC is also known for its powerful, beneficial behavioural consequences and its modulating effects on neurobiology and brain ultrastructure (van Praag *et al*. 2000; Nithianantharajah and Hannan 2006). For example, EC improves learning and memory (Simpson and Kelly 2011), reduces anxiety (Reichmann *et al*. 2016), promotes stress resilience (Crofton *et al*. 2015) and ameliorates disease phenotypes in animal models of neuropsychiatric disease (Pang and Hannan 2013). In contrast, there is also evidence that increased EC could have negative effects on laboratory animals. Specifically, it has been shown that EC induces agonistic encounters and increases aggressive behaviour in male mice, leading to increased anxiety-like behaviour, higher corticosterone levels and eventually to physical and psychological injuries (Cabrera-Muñoz etal., 2022; McQuaid et al., 2018; Sowndharya & Rajan, 2024). EC has also been associated with non-adaptive behaviours in rodents as well as with enhanced alcohol intake (Berardo *et al*. 2016; González-Pardo *et al*. 2019; Suárez *et al*. 2020; Corredor *et al*. 2022). In addition, it has been found that EC can lead to chronic stress (Moncek *et al*. 2004; Domínguez-Oliva *et al*. 2025) and behavioural despair (Guven *et al*. 2022).

In a study investigating the effects of EC on stress-induced neuronal activation, we have previously shown that colitis modulates the effects of EC on stress-induced neural activity, indicating an interaction between EC and gut-brain axis signalling (Reichmann et al., 2013). In addition, we and others found that EC can enhance peripheral inflammation and increase immune system activity, suggesting that EC might have negative effects in animal models of inflammatory diseases (Brod et al., 2017; Reichmann et al., 2013). In this study, we therefore set out to systematically investigate, if EC exerts positive or negative effects on MGB axis signalling in an animal model of UC. We investigated how EC affects gut-brain axis signalling in the course of DSS colitis, and if EC is able to modify colitis-induced changes to behaviour and neurobiology. To gain mechanistic insights, we characterized colitis and EC effects on all levels of the MGB axis by assessing inflammatory processes, intestinal dysbiosis and metabolic shifts.

## Results

### Enhanced environmental complexity (EC) exacerbates DSS-induced colonic inflammation

To analyse if EC modulates MGB axis signalling in the course of colitis, we first investigated the effects of DSS treatment after 9 weeks of EC housing on disease activity and colonic inflammation in male and female mice (Fig. 1A; Suppl. Fig. 1). We found that female mice kept in standard environment (SE) and treated with DSS for 7 days lost weight at the end of the treatment period (day 8-10) compared with baseline (day 1), which partially recovered after stopping DSS treatment (day 11-12; Fig. 1B). Interestingly, however, this body-weight loss was much more pronounced in female DSS-treated EC mice than in SE-housed controls (time x group interaction: F_(21, 279)_ = 25.550; P<0.001), and these animals did not regain any of the lost weight after stopping DSS (Fig. 1B). This finding was not the result of increased DSS intake in the EC group, given that the amount of liquid consumed during the treatment period was similar between the two colitis groups (Suppl. Fig. 2A). DSS treatment also significantly shortened the colon of female mice (DSS main effect: F_(1, 33)_ = 43.050; P<0.001; Fig. 1C), and while female EC/DSS animals had on average the shortest colons (5.890 ± 0.211 cm), this failed to reach statistical significance compared to SE/DSS mice (6.440 ± 0.197 cm; P = 0.184). As expected, DSS treatment increased the disease activity index (DAI; main effect DSS: F_(1, 34)_ = 81.100; P<0.001) compared to non-treated mice, but female EC/DSS mice had higher DAI scores than SE/DSS mice both at the end of DSS treatment (day 8; DSS x housing interaction: F_(1, 33)_ = 6.750; P=0.014; Fig. 1D) and five days later (day 12; DSS x housing interaction: F_(1, 34)_ = 8.264; P=0.007; Suppl. Fig. 2B). Additionally, DSS treatment significantly reduced spleen weight in female EC/DSS mice compared to controls, but also SE/DSS mice (day 9; DSS x housing interaction: F_(1, 36)_ = 10.460; P=0.003; Fig. 1E). We also detected an increase of colonic myeloperoxidase (MPO) content, a marker of leukocyte infiltration, in the colons of female DSS-treated mice (day 9; DSS main effect: F_(1, 33)_ = 35.510; P<0.001), again significantly more pronounced in the EC/DSS group (DSS x housing interaction: F_(1, 33)_ = 7.593; P=0.0100; Fig. 1F).This result is also reflected at the histological level. The colons of female DSS-treated mice were characterized by extensive inflammatory cell infiltration, crypt loss and goblet cell loss, effects not observed in healthy controls (DSS main effect: F_(1, 28)_ = 353.400; P<0.001; Fig. 1G-H). However, inflammation severity and tissue remodelling were again more pronounced in the EC/DSS group than the SE/DSS group (DSS x housing interaction: F_(1, 28)_ = 6.087; P=0.020; Fig. 1G-H). In male mice, DSS treatment reduced body weight (time x group interaction: F_(5.459, 65.510)_ = 7.855; P<0.001), colon length (F_(1, 36)_ = 45.730; P<0.001) and spleen weight (F_(1, 36)_ = 5.177; P=0.029) and increased the DAI (F_(1, 36)_ = 50.480; P<0.001; Suppl. Fig. 1). However, in contrast to female mice, EC had no modifying effect on these parameters (Suppl. Fig. 1).

**Figure 1.**
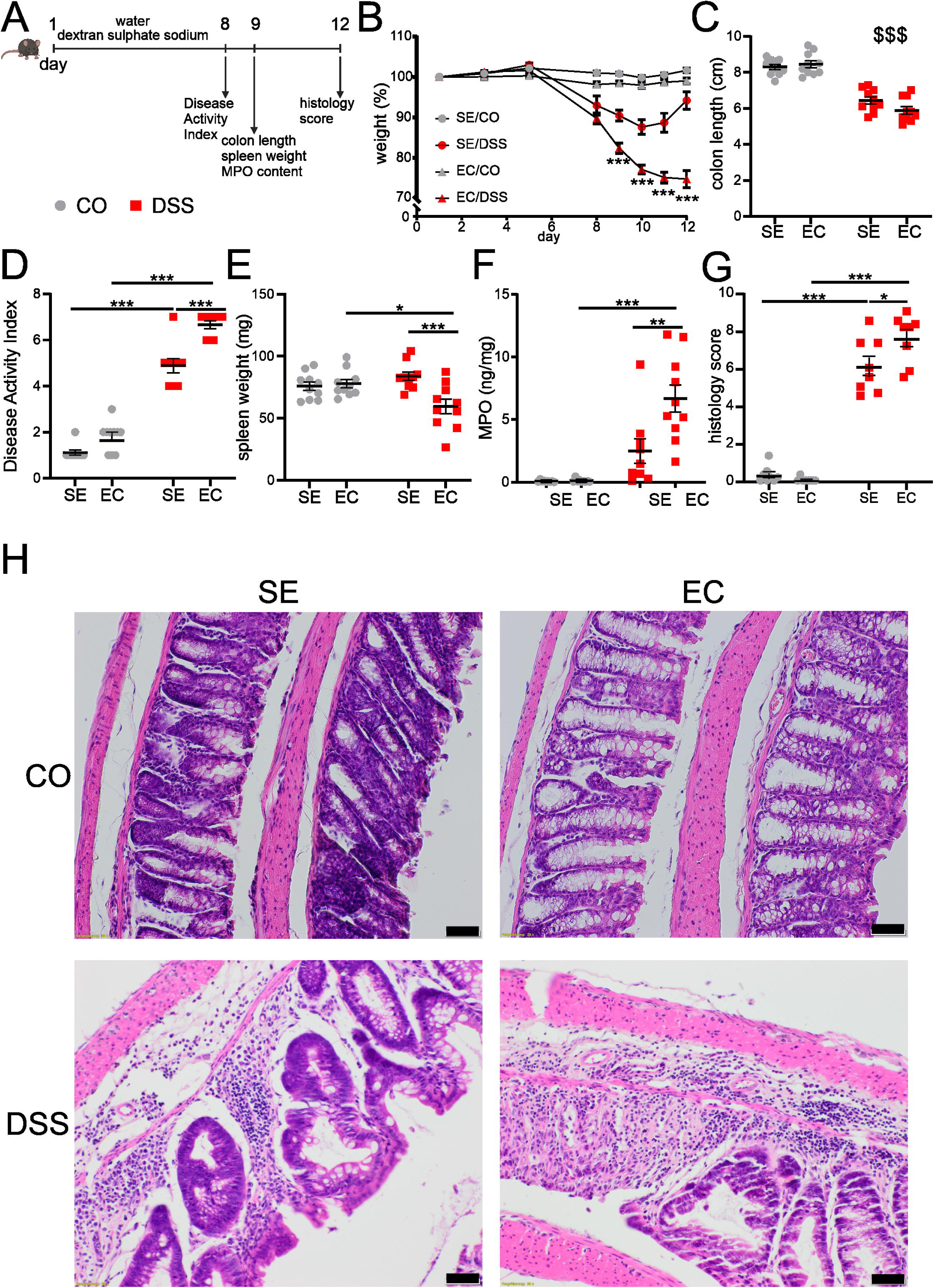
Enhanced environmental complexity exacerbates colitis-induced disease phenotypes in female mice. (A) Experimental timeline of dextran sulphate sodium (DSS) treatment and readouts. (B) Effects of enhanced environmental complexity (EC) and DSS treatment on body weight over time. Repeated-measures two-way ANOVA. ***p < 0.001 EC/DSS vs SE/DSS. n = 9-10/group. Effects of EC and DSS treatment on (C) colon length, (D) disease activity index, (E) spleen weight, (F) colonic myeloperoxidase (MPO) levels and (G) histology score. Data are presented as mean ± SEM. Two-way ANOVA followed by Tukey post-hoc test in case of a significant treatment X housing interaction. main effect of DSS: ^$$$^p < 0.001. Tukey post-hoc testing: ***p < 0.001, **p < 0.01, *p < 0.05; n = 9-10/group. (H) Representative images of colonic tissues collected 5 days after stopping DSS treatment (day 12) stained with Hematoxylin and Eosin. scalebar: 50 µm. CO = control; DSS = dextran sulphate sodium; EC = enhanced environmental complexity; SE = standard environment.

### Colitis and EC exert distinct effects on immune cell populations along the gut-brain axis

Previous studies found that EC is able to alter the activity of the immune system (Brod *et al*. 2017; Xiao *et al*. 2019). Therefore, we investigated whether the more severe disease course of female DSS-treated mice housed in EC is a consequence of enhanced colonic immune cell infiltration (flow cytometry gating strategy in Suppl. Fig. 3). We found that innate immune cells including monocytes, macrophages, eosinophils and neutrophils are more abundant in the colon of DSS-treated mice compared to non-treated controls independent of housing conditions (Fig. 2A-D; DSS main effect: F_(1, 35)_ = 61.130; P<0.001 for monocytes; F_(1, 35)_ = 13.950; P<0.001 for macrophages; F_(1, 35)_ = 21.750; P<0.001 for eosinophils; F_(1, 35)_ = 25.350; P<0.001 for neutrophils). In addition, we detected higher numbers of T helper, cytotoxic T cells and natural killer (NK) cells in DSS-treated mice (Fig. 2E-G; DSS main effect: F_(1, 15)_ = 9.898; P=0.007 for T helper cells; F_(1, 15)_ = 10.900; P=0.005 for cytotoxic T cells; F_(1, 35)_ = 6.727; P=0.014 for NK cells), but no groups difference in NK T cell numbers (Fig. 2H). EC had no effect on any of these cell populations in healthy animals and it also did not modify the DSS-induced colonic cell infiltration. This suggests that the more pronounced weight loss and the higher disease activity of EC mice with colitis are not caused by an enhanced number of immune cells in colonic tissue.

**Figure 2.**
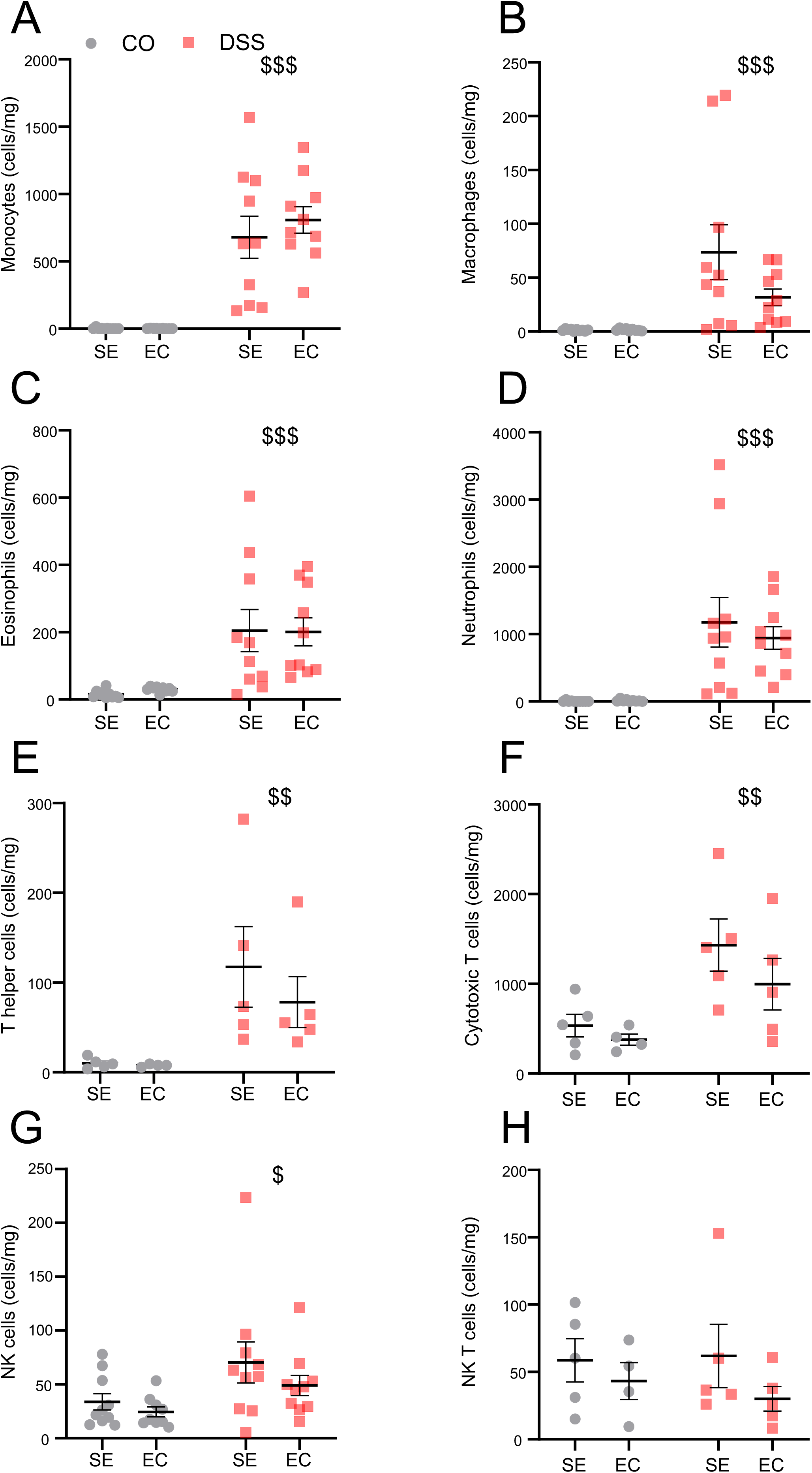
Colitis enhances colonic immune cell infiltration independent of housing condition. Flow cytometric quantification of immune cells expressed as number of cells per mg of colonic tissue using the gating strategy described in Suppl. Fig. 3. (A) monocytes; (B) macrophages; (C) eosinophils; (D) neutrophils; (E) T helper cells; (F) cytotoxic T cells; (G) NK cell and (H) NK T cells. CO = control; DSS = dextran sulphate sodium; EC = enhanced environmental complexity; SE = standard environment. Data are presented as mean ± SEM. Two-way ANOVA. main effect of DSS: ^$$$^p < 0.001, ^$$^p < 0.01, ^$^p < 0.05. n = 4-10/group.

DSS colitis is known to lead to systemic immune responses (Hall *et al*. 2011) and thus we also assessed how EC and DSS colitis affect the number of circulating immune cells. Similar to the colon, we found that DSS treatment leads to an increased number of monocytes, macrophages and neutrophils in the blood circulation, an effect independent of housing condition (Fig. 3A, B, D; DSS main effect: F_(1, 35)_ = 56.640; P<0.001 for monocytes; F_(1, 35)_ = 51.510; P<0.001 for macrophages; F_(1, 35)_ = 45.270; P<0.001 for neutrophils). However, in contrast to the colon, the percentage of circulating eosinophils was significantly lower after DSS treatment under both housing conditions (DSS main effect: F_(1, 35)_ = 17.990; P<0.001; Fig. 3C), which might reflect an increase in pro-apoptotic signals of these cells or an enhanced recruitment to the site of inflammation. We observed no differences between the groups in cells related to adaptive immunity including T helper cells, cytotoxic T cells, NK cells and NK T cells (Fig. 3E-H). Like in the colon, EC did not modify the number of any measured immune cell population both in healthy animals and animals with colitis.

**Figure 3.**
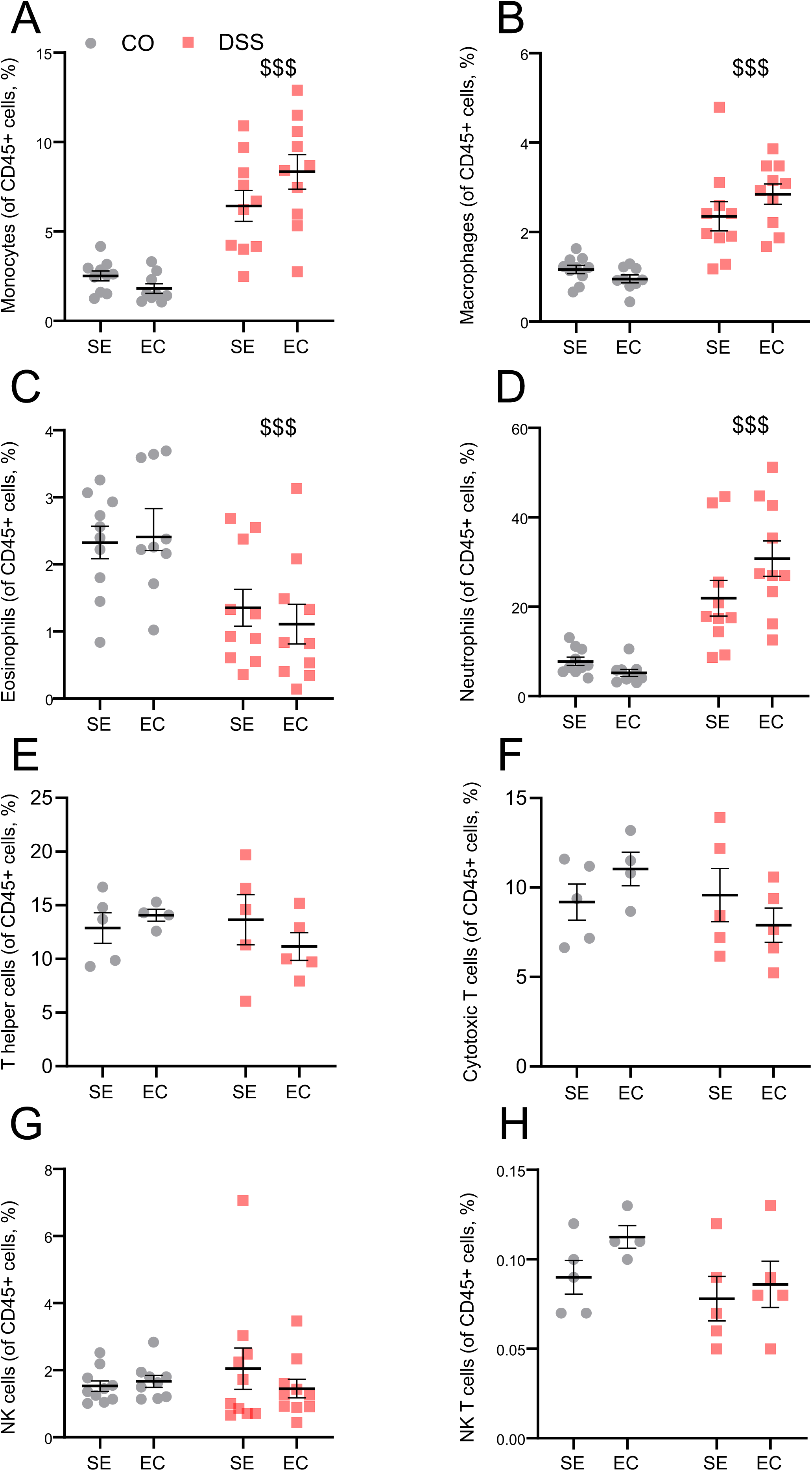
Colitis alters the proportion of circulating innate immune cells independent of housing condition. Flow cytometric quantification of circulating immune cells expressed as % of CD45^+^ cells using the gating strategy described in Suppl. Fig. 3. (A) monocytes; (B) macrophages; (C) eosinophils; (D) neutrophils; (E) T helper cells; (F) cytotoxic T cells; (G) NK cells and (H) NK T cells. CO = control; DSS = dextran sulphate sodium; EC = enhanced environmental complexity; SE = standard environment. Data are presented as mean ± SEM. Two-way ANOVA. main effect of DSS: ^$$$^p < 0.001. n = 4-10.

We and others have previously shown that the systemic inflammation during DSS colitis reaches distant organs including the brain (Sroor *et al*. 2019; Gampierakis *et al*. 2021) and thus we also investigated immune cell populations in brain tissue (flow cytometry gating strategy in Suppl. Fig. 4). Analyses revealed that, among innate immune cells, colitis led to an increase in monocyte-derived macrophages, but did not alter the number of monocytes, eosinophils or neutrophils (Fig. 4A-D; DSS main effect: F_(1, 35)_ = 4.665; P=0.038). We also observed a lower number of cytotoxic T cells after DSS colitis (DSS main effect: F_(1, 15)_ = 5.304; P=0.036), while other CNS T cell populations, as well as NK cells, were not affected by colitis (Fig.4E-H). In contrast to immune cell populations in the colon and the circulation, EC had notable effects on immune cell subpopulations in the brain. Independent of DSS treatment, EC-housed mice had lower brain neutrophil cell counts compared to mice kept in SE (Fig. 4D; EC main effect: F_(1, 35)_ = 6.042; P=0.019) and a lower number of T helper cells (Fig. 4E; EC main effect: F_(1, 15)_ = 7.657; P=0.014), indicating reduced immune cell infiltration via the blood-brain-barrier. Finally, EC also reduced the number of activated microglial cells when compared to the SE-housed counterparts (EC main effect: F_(1, 35)_ = 4.446; P=0.042; Fig. 4I), which might indicate a reduced activity of the brain’s immune defence.

**Figure 4.**
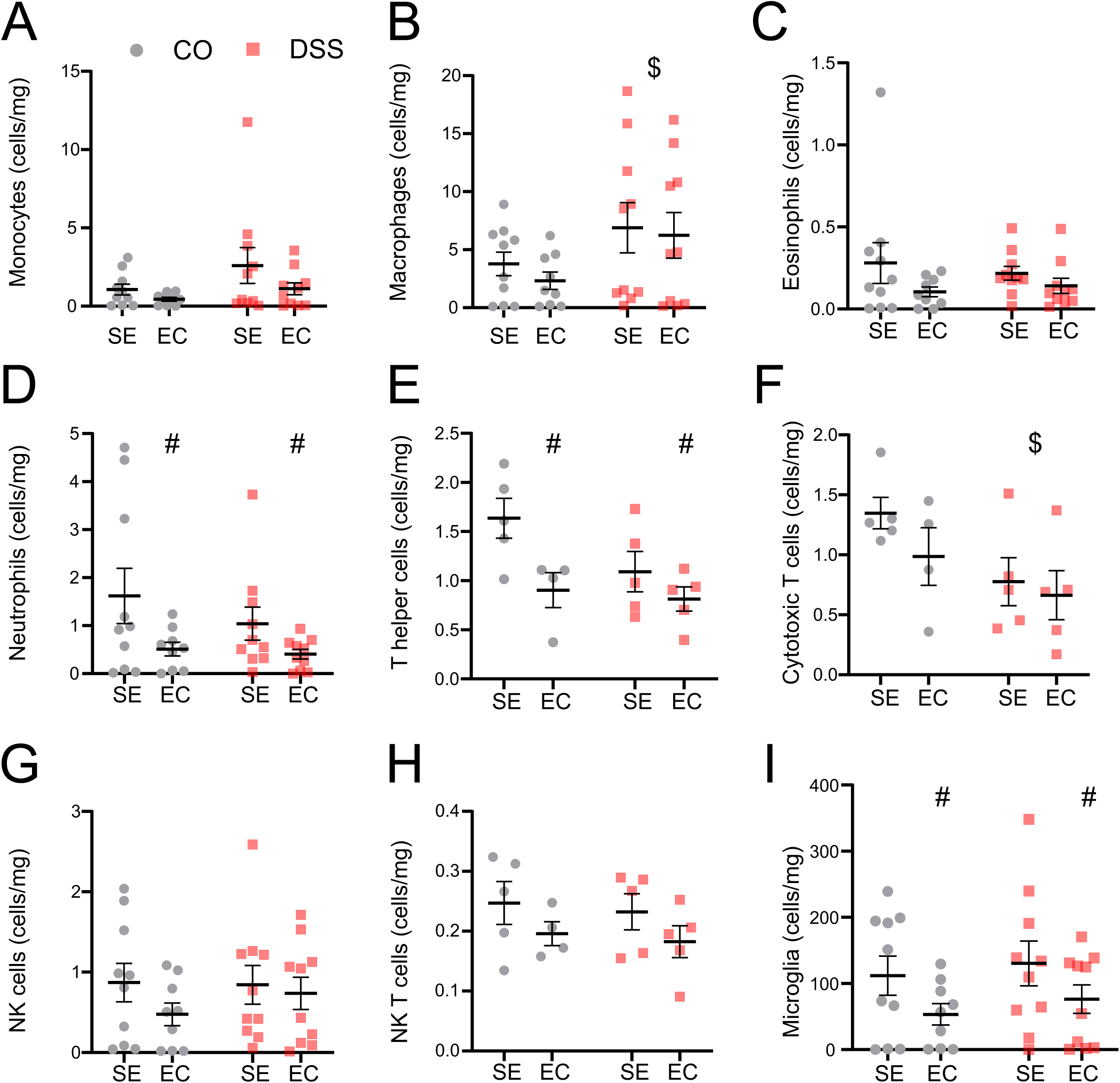
Effects of colitis and enhanced environmental complexity on immune cell subpopulations within the brain. Flow cytometric quantification of immune cell populations expressed as number of cells per mg of brain tissue using the gating strategy described in Suppl. Fig. 4 (A) monocytes; (B) macrophages; (C) eosinophils; (D) neutrophils; (E) T helper cells; (F) cytotoxic T cells; (G) NK cells,(H) NK T cells and (I) microglial cells. CO = control; DSS = dextran sulphate sodium; EC = enhanced environmental complexity; SE = standard environment. Data are presented as mean ± SEM. Two-way ANOVA. main effect of DSS: ^$^p < 0.05. main effect of EC: ^#^p < 0.05. n = 4-10/group.

### EC heightens colitis-induced metabolite alterations across the gut-brain axis

Given that female EC/DSS mice did not show changes in the number of circulating or tissue-resident immune cells compared to SE/DSS mice, we next investigated effects on inflammatory mediators and metabolites. To do this, we first examined circulating corticosterone levels to assess the stress levels of the animals. We found that both colitis (DSS main effect: F_(1, 35)_ = 18.730; P<0.001) and EC (EC main effect: F_(1, 35)_ = 16.690; P<0.001) increased corticosterone (Fig. 5A), but no treatment X housing interaction, suggesting that both interventions increase stress levels independent of each other. However, planned comparisons revealed that mice in the EC/DSS group had the highest corticosterone levels (80.893 ± 14.804), which were significantly higher than in the SE/DSS group (37.310 ± 11.825; P=0.001; Fig. 5A). This indicates a synergistic negative impact of both interventions on the stress axis. In addition to that, we also examined pro-inflammatory cytokine levels in the blood circulation, as potential mediators of the negative consequences of EC during colitis (Fig. 5B-F). While we detected a housing condition-independent rise of IL-18 (Fig. 5B; DSS main effect: F_(1, 35)_ = 24.230; P < 0.001), TNF-α (Fig. 5C; DSS main effect: F_(1, 36)_ = 19.960; P < 0.001) and GRO-α (Fig. 5D; DSS main effect: F_(1, 36)_ = 43.200; P<0.001) after DSS treatment, other cytokines showed EC-induced modulations. Specifically, we observed a decrease of IL-22 in EC-housed animals with and without colitis (Fig. 5E; EC main effect: F_(1, 36)_ = 4.177; P=0.048) and a rise of circulating IL-6 levels (Fig. 5F; EC main effect: F_(1,36)_ = 8.073; P=0.007). Experimental colitis also increased IL-6 levels (Fig. 5F; DSS main effect: F_(1,36)_ = 11.320; P=0.002) without a significant interaction between the two factors. Interestingly, however, planned comparisons of the two colitis groups revealed a trend towards higher levels in EC/DSS mice compared to SE/DSS mice (P=0.075).

**Figure 5.**
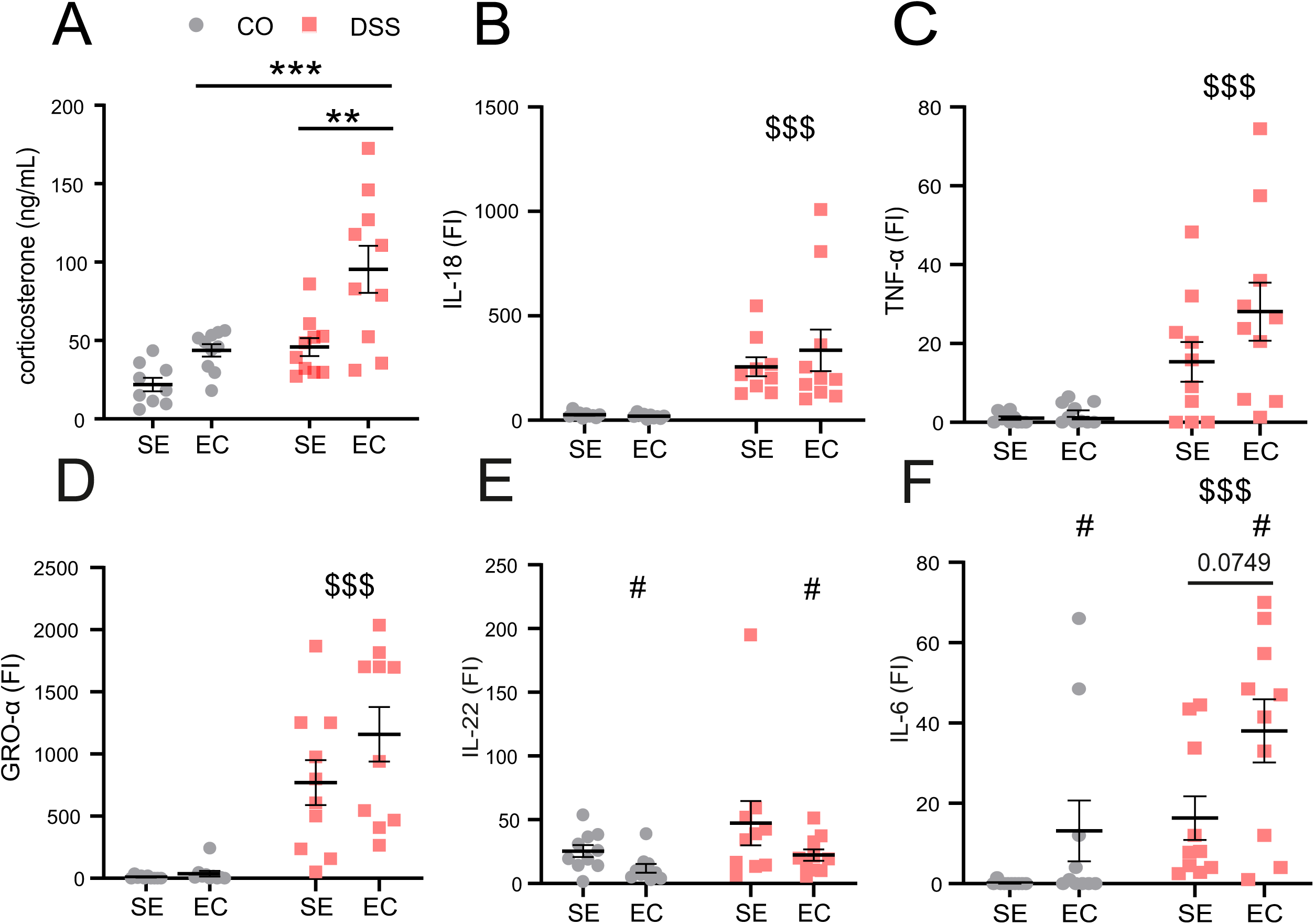
Colitis and enhanced environmental complexity alter circulating corticosterone and cytokines levels. Effects of colitis and enhanced environmental complexity (EC) on plasma levels of (A) corticosterone (ng/mL) measured with corticosterone ELISA Kit; (B) interleukin (IL)-18; (C) TNF-α; (D) GRO-α; (E) IL-22 and (F) IL-6 expressed as fluorescence intensity (FI) minus background measured with Procartaplex Multiplex ELISA Assay. CO = control; DSS = dextran sulphate sodium; EC = enhanced environmental complexity; SE = standard environment. Data are presented as mean ± SEM. Two-way ANOVA followed by Tukey post-hoc testing. main effect of DSS: ^$$$^p < 0.0001. main effect of EC: ^#^p < 0.05. Tukey post-hoc testing: ***p < 0.001, **p < 0.01. n = 10/group.

Next, we explored whether EC and/or colitis affect metabolic profiles along the gut-brain axis of female mice by performing plasma, brain tissue, cerebrospinal fluid (CSF) and brain interstitial fluid (ISF) metabolomics. After applying analytical-quality criteria, the detected metabolite numbers were 330 in plasma, 356 in brain tissue, 275 in CSF, and 286 in ISF samples. Detailed analysis revealed that EC and colitis induced numerous significant metabolite alterations in all sample types investigated (Suppl. Table 1). Based on absolute numbers, we found the largest metabolic changes in the CSF of healthy mice kept in EC (EC/CO). Compared to SE-housed controls (SE/CO), 111 metabolites were more abundant in the EC/CO group (Fig. 6A; Suppl. Table 2), suggesting that EC strongly increases the secretion of various metabolites from the brain into the CSF and/or the reabsorption of metabolites from the CSF. Even more metabolite changes (123 significantly different metabolites) were detected when comparing EC/CO with EC/DSS. The vast majority (121) of those metabolites were again more abundant in the EC/CO group (Fig. 6B; Suppl. Table 2) suggesting that the CSF metabolome of healthy EC mice is very different from the metabolome of all other investigated groups. Given that the CSF metabolome of the EC/DSS group is much more similar to the SE/CO group than the EC/CO group, it appears that colitis disrupts the ability of EC to alter CSF metabolite dynamics. Supporting this view, we found a high overlap of 98 metabolites altered both in the EC/CO vs. SE/CO comparison and also the EC/CO vs. EC/DSS comparison (Suppl. Fig.5A). Similar findings were made in the plasma of the EC/CO group (Suppl. Table 3), where the vast majority of significantly different metabolites had again higher concentrations compared to SE/CO (45 out of 50 significantly different metabolites) and EC/DSS (53 out of 63 metabolites). Most of the metabolites altered between EC/CO and SE/CO belong to the class of amino acids and derivatives or to energy metabolism metabolites (Fig. 6C; Suppl. Table 3), which might reflect the exercise component of EC that can have effects on multiple organ systems thereby changing the plasma metabolome. Interestingly, experimental colitis in EC-housed animals led to a decrease in metabolite concentrations spanning most major metabolite classes compared to EC/CO (Fig.6D; Suppl. Table 3), but in contrast to the CSF metabolome, we observed much less overlap to the metabolite alterations between EC/CO and SE/CO (Suppl. Fig.5B). While the brain ISF showed comparatively fewer metabolic alterations between the two housing conditions (Suppl. Fig.5C; Suppl. Table 4), EC, as expected, had strong effects on the brain metabolome of healthy animals (48 significantly different metabolites), especially regarding free fatty acids levels (Fig.6E; Suppl. Table 5). A similar number of metabolites was altered between EC/CO and EC/DSS mice (Fig.6F; 51 significantly different metabolites) suggesting again a disruption of the metabolic effects of EC by experimental colitis. Interestingly, some of the metabolites changed by EC in healthy mice were altered in more than one compartment investigated (Fig.6G). The most pronounced alteration in this regard is tiglylcarnitine, which was changed in all compartments investigated. Specifically, it was upregulated in the CSF and in the plasma, but downregulated in the ISF and brain of EC-housed animals (Fig.6A,C,E; Suppl. Fig.4C). Tiglylcarnitine is a short branched-chain acylcarnitine, which is involved in branched-chain amino acid catabolism and beta-oxidation (Dambrova *et al*. 2022), thus suggesting strong effects of EC on energy metabolism throughout the body. Three metabolites were changed in three out of 4 compartments investigated, including the nucleotide base uracil that was upregulated in the brain, plasma and CSF. In addition, 2-ketobutyric acid, a short-chain keto acid involved in the catabolism of amino acids, was upregulated in the plasma and CSF, while it was downregulated in the brain after EC. Finally, the free fatty acids FFA C18:3 were downregulated in the plasma, while they were upregulated in the brain and CSF. Fatty acids with 18 carbon atoms and 3 double bonds are polyunsaturated fatty acids (PUFAs) including α-Linolenic acid (ALA) and γ-Linolenic acid (GLA). The most common form is ALA, an essential omega-3 PUFA with important roles in cell signalling, metabolism, and inflammation regulation (Yuan *et al*. 2022). A decrease of ALA in the plasma, no change in brain ISF, but higher levels in brain tissue and CSF suggest higher peripheral utilization or clearance of ALA, but possible accumulation in the brain cells after EC. 41 metabolites were altered in 2 compartments after EC (Fig. 6G, Suppl. Table 6), in most cases in plasma and CSF (25 metabolites) or in brain and CSF (13 metabolites). This might indicate strong signalling from the periphery to the CNS during EC and altered brain metabolism due to EC, respectively.

**Figure 6.**
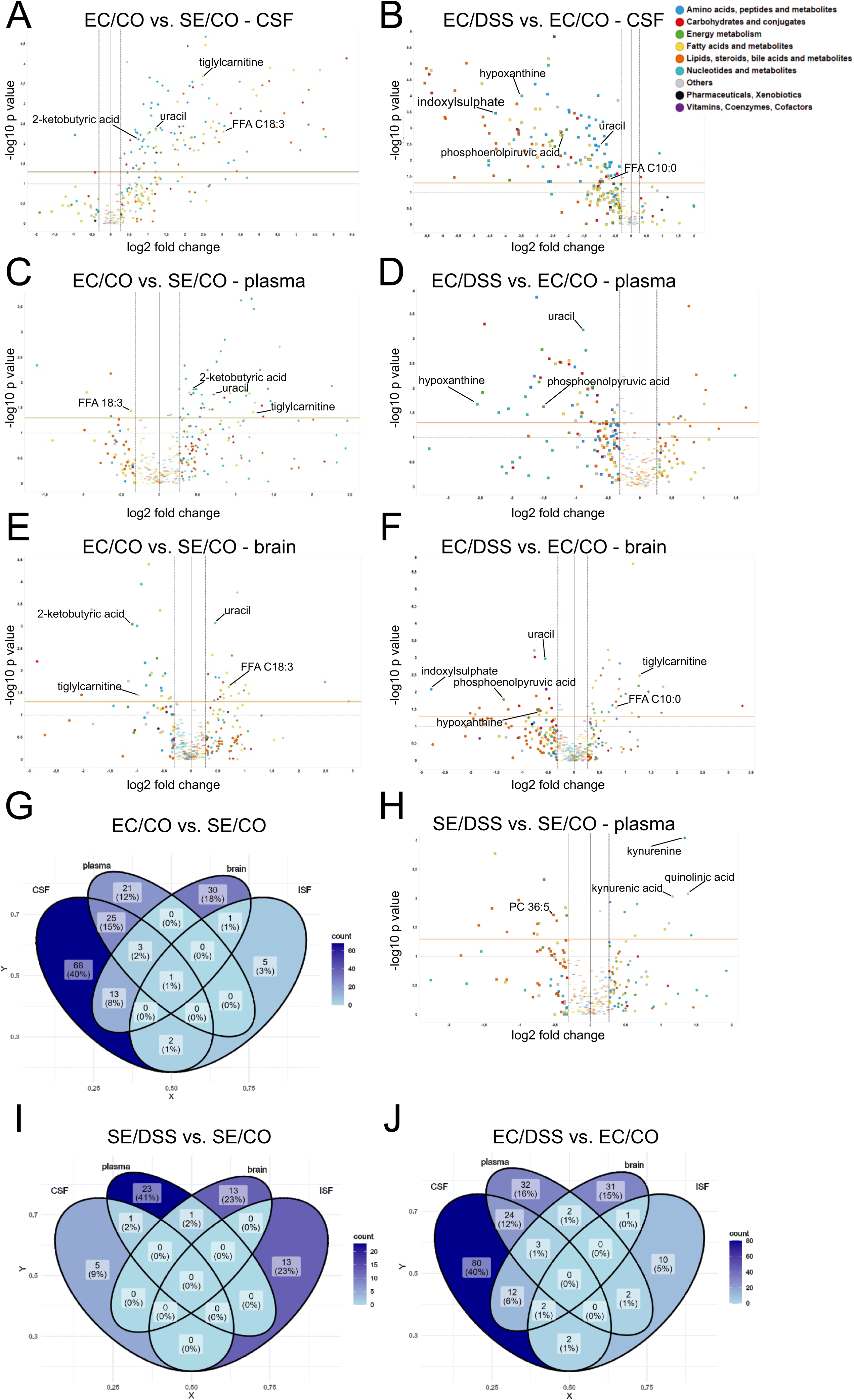
Effects of enhanced environmental complexity (EC) and colitis on brain tissue, brain interstitial fluid, cerebrospinal fluid and plasma metabolome. Volcano plots displaying statistically significant altered metabolites between EC/CO and SE/CO mice in (A) cerebrospinal fluid (CSF), (C) plasma and (E) brain tissue. Volcano plots displaying statistically significant altered metabolites between EC/DSS and EC/CO mice in (B) CSF, (D) plasma and (F) brain tissue. Venn diagrams highlighting the overlap of significantly altered metabolites between all 4 compartments investigated for the comparison (G) EC/CO vs. SE/CO, (I) SE/DSS vs. SE/CO and (J) EC/DSS vs. EC/CO. (H) Volcano plot displaying significantly altered metabolites in plasma of SE/DSS and SE/CO mice. CO = control; DSS = dextran sulphate sodium; EC = enhanced environmental complexity; SE = standard environment.

Colitis also led to significant metabolic alterations in all compartments investigated. Interestingly, these effects were much more pronounced under EC conditions (Suppl. Table 1). When analysing the effects of colitis under standard conditions (SE/DSS vs. SE/CO), most changes were detected in the plasma (Fig.6H; Suppl. Table 3; 25 significantly different metabolites), indicating metabolites altered in the systemic circulation in response to peripheral inflammation. For example, for phosphatidylcholines – which have been linked to reduced tissue damage and lower inflammation (Wen *et al*. 2023) – we detected lower levels in mice with colitis. In addition, kynurenine and its metabolites kynurenic acid and quinolinic acid had higher levels in the colitis group indicating a disruption of tryptophan metabolism (Sheibani *et al*. 2023). Notably, much more plasma metabolites were altered due to colitis under EC conditions (DSS/EC vs. CO/EC; 63 significantly metabolites, see Fig. 6D) suggesting that EC facilitates colitis-induced plasma metabolite changes. DSS-treated animals housed in SE showed also significantly increased concentrations of 7 metabolites in their brain, 1 in their ISF and 1 in their CSF and decreased concentrations of 7 compounds in the brain, 12 in the ISF and 5 in the CSF compared to their respective controls (Suppl. Fig. 5D-F). This is again significantly less than under EC conditions, where we observed 51 altered brain metabolites, 17 altered ISF metabolites and 123 altered CSF metabolites (Suppl. Table1, Fig. 6B, F, Suppl. Fig. 5G) Only few colitis-induced metabolite alterations under standard housing conditions occurred in multiple compartments (Fig. 6I). One metabolite – kynurenine – was upregulated in brain and plasma, while PC 36:5 was downregulated both in the CSF and in the plasma in mice with colitis compared with non-treated controls (Fig. 6H, Suppl. Fig. 5D, F). This is again significantly less than under EC conditions, where we detected 43 metabolites that were changed in 2 compartments, mainly in plasma and CSF (24 metabolites) or brain and CSF (12 metabolites; Suppl. table 7) and 5 metabolites changed in 3 compartments (Fig. 6J). Among those changed in 3 compartments, hypoxanthine was significantly decreased in plasma, brain and CSF in the EC/DSS group, as well as uracil and phosphoenolpyruvic acid. Moreover, capric acid (FFA C10:0) and tiglylcarnitine were upregulated in the brain and the ISF of the EC/DSS group, while they were downregulated in the CSF, compared with non-treated controls (Fig 6 B,D,F, Suppl. Fig. 5G).

When directly comparing the metabolome of DSS-treated animals under standard and enriched housing conditions (EC/DSS vs. SE/DSS), we also detected numerous differences in all compartments investigated (Suppl. Table 1). Specifically, in the EC/DSS group 9 metabolites were less abundant in plasma, 12 in the brain, 5 in the ISF and 3 in the CSF than in the SE/DSS group. In addition, DSS-treated EC-housed mice had significantly higher levels of 16 plasma metabolites, 7 brain metabolites, 5 ISF metabolites and 1 CSF metabolite (Fig. 7A-D; Suppl. Table 2-5). Interestingly, the strongest upregulated metabolites in plasma and brain of the EC/DSS group are microbiota-derived metabolites such as trimethylamine-N-oxide (TMAO) and putrescine as well as primary and secondary bile acids including muricholic acid/cholic acid/ursocholic acid, allocholic acid, 7-ketochenodeoxycholic acid and deoxycholic acid. Like TMAO, secondary bile acids are generated by gastrointestinal microbiota (Winston and Theriot 2020) suggesting that EC altered the gastrointestinal community composition during colitis. Few of these metabolites were changed in multiple compartments (Fig. 7E), but phenylacetic acid, another microbiota-derived metabolite, was downregulated in both ISF and CSF, gluconic acid was downregulated in plasma and brain and deoxycholic acid was upregulated in plasma and brain of the EC/DSS group (Fig. 7A-D). To investigate the metabolite alterations between SE/DSS and EC/DSS in more detail, we also performed pathway analysis to identify relevant underlying biological processes. In the plasma metabolome this analysis revealed an enrichment of multiple pathways related to various biological functions such as cell death, digestive system, endocrine system, immune system, lipid metabolism, nervous system, signal transduction and transport and catabolism in the EC/DSS group (Fig. 7F). Metabolites driving these findings are mainly the lipid metabolites PC 32:2, PC 33:2/PE 36:2, PC 34:4, PC 36:4, PC 38:4, Lyso PC 18:3, Lyso PC 20:4, Lyso PE 18:0 (Suppl. Table 8), which are not only involved in lipid metabolism, but also, to a varying extent, in immune system activity. We also detected numerous pathway alterations in the brain metabolome between SE/DSS and EC/DSS, which are related to a wide range of processes such as aging, amino acid metabolism, carbohydrate metabolism, cell growth and death, the endocrine system, energy metabolism, glycan biosynthesis and metabolism, the immune system, lipid metabolism, metabolism of co-factors and vitamins, metabolism of other amino acids, the nervous system, nucleotide metabolism, signal transduction and transport and catabolism (Fig. 7G). Of particular interest among the changed pathways of the brain metabolome are the pathways related to the nervous system pathway group reflecting colitis-induced changes to neurobiology that are modified by EC. Specifically, we found significant alterations in synaptic signalling, retrograde endocannabinoid signalling and the long-term depression (LTD) pathways in the EC/DSS group (Suppl. Table 9). The latter two processes are directly connected through the mechanism of endocannabinoid-mediated synaptic plasticity (Robbe *et al*. 2002) suggesting that EC might alter the capacity for neuronal remodelling during colitis. Like in the plasma, we also found differences in immune system-related pathways in the brain metabolome. However, in contrast to the plasma metabolome, all the significantly altered immune system pathways are downregulated in the brain metabolome of the EC/DSS group, because of the less abundant energy metabolites adenosine diphosphate (ADP), dihydroxyacetone phosphate (DHAP), nicotinamide adenine dinucleotide phosphate (NADP), flavin adenine dinucleotide (FAD) involved in the detected immunological processes (Suppl. Table 10). This suggests that EC, although exacerbating peripheral inflammation in mice with colitis, prevents excessive neuroinflammation in our model, which is in line with the reduced brain T cell counts after EC detected in this study and other reports that observed reduced neuroinflammation after EC in multiple disease models (Gonçalves *et al*. 2026). In contrast to plasma and brain, pathway analysis of the CSF and ISF compartments did not reveal significantly altered pathways between SE/DSS and EC/DSS. This might be related to the small number of differentially abundant metabolites in these compartments.

**Figure 7.**
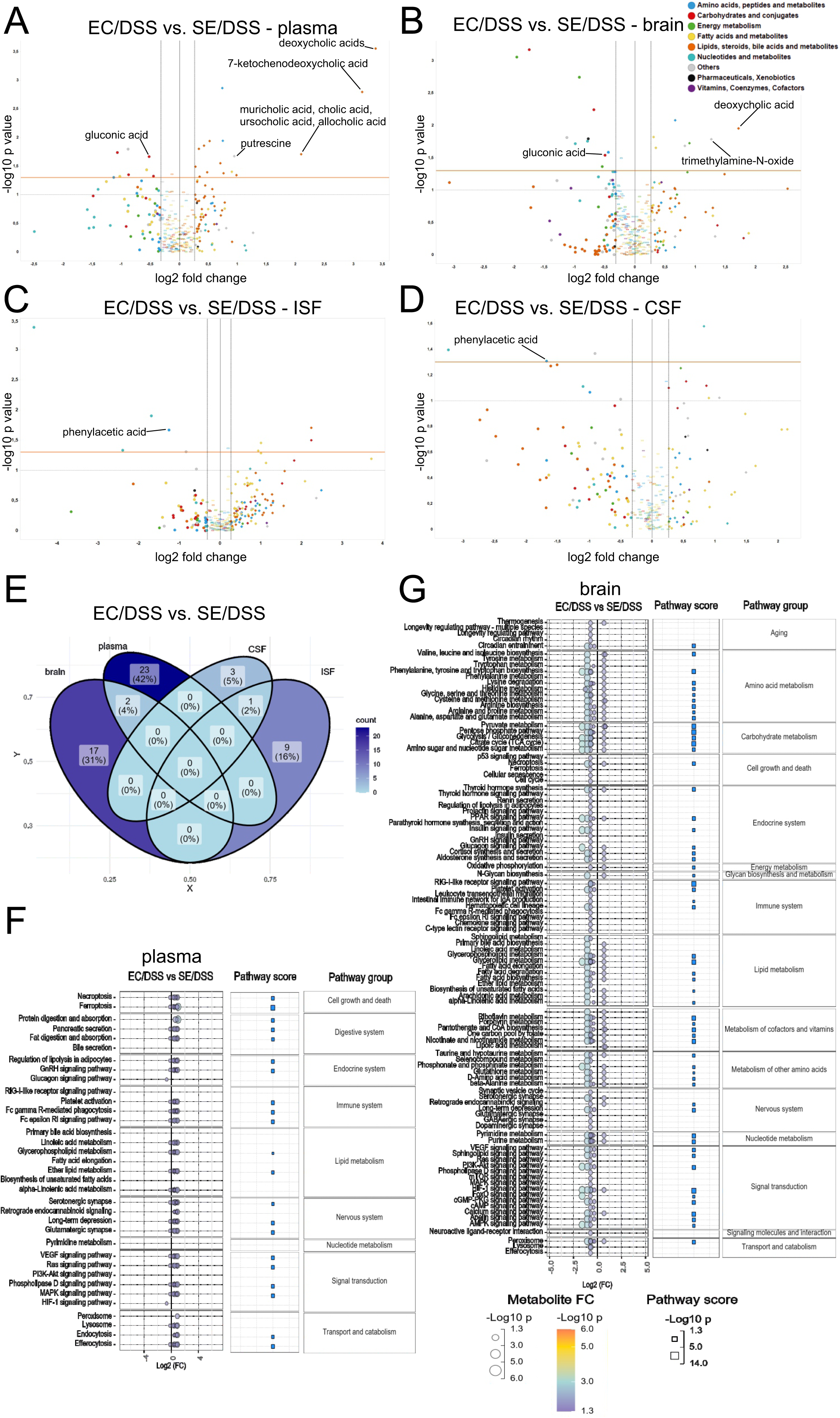
Enhanced environmental complexity (EC) modulates colitis-induced metabolome changes across the gut-brain axis. Volcano plots displaying significantly altered metabolites between EC/DSS and SE/DSS mice in (A) plasma, (B) brain tissue, (C) brain interstitial fluid (ISF) and (D) cerebrospinal fluid (CSF). (E) Venn diagram highlighting the overlap of significantly altered metabolites between all 4 compartments investigated for the comparison EC/DSS vs. SE/DSS. KEGG pathway analysis of significantly altered metabolites in (F) plasma and (G) brain tissue for the comparison EC/DSS vs. SE/DSS. CO = control; DSS = dextran sulphate sodium; EC = enhanced environmental complexity; SE = standard environment

### EC modifies colitis-induced changes to the gastrointestinal microbiome

Given that metabolomics revealed microbiota-derived metabolites as the strongest downregulated metabolites in the EC/DSS group compared to the SE/DSS group, we next assessed differences in microbial community composition after EC and/or colitis using 16S sequencing. Richness (Fig. 8A) and evenness (Fig. 8B) were both significantly reduced in mice with colitis, independent of housing condition; similarly, the Shannon index (Fig. 8C) showed a significant reduction in alpha diversity, also independent of housing. These findings indicate a dysbiotic shift in the gut microbiota, characterized by loss of diversity and selective enrichment of inflammation-adapted taxa. We also detected a strong difference in beta diversity between mice with and without colitis using principal coordinates analysis (PCoA) based on Bray-Curtis dissimilarity (Fig. 8D; R² = 0.412, P<0.001), but did not observe clear separate clusters for the different housing conditions. Taxonomic composition analysis also showed large differences in relative abundance in response to colitis, while EC had less pronounced effects on bacterial taxonomic composition. At phylum level, we found that the most dominant taxa in healthy animals are Firmicutes and Bacteriodota. In mice with colitis that is also the case, but we also observed a considerable increase in Verrucomicrobiota and Cyanobacteria, compared to untreated groups (Fig. 8E). At genus level, colitis also led to major shifts in microbial composition, while the most abundant genera were not altered by EC in healthy mice (Fig. 8F). Specifically*, Akkermansia, Alistipes,* Clostridia_UCG-014, Clostridium sensu stricto_1 and *Turicibacter* were more abundant in DSS-treated groups, while *Lactobacillus* and *Alloprevotella* were less abundant in animals with colitis.

**Figure 8.**
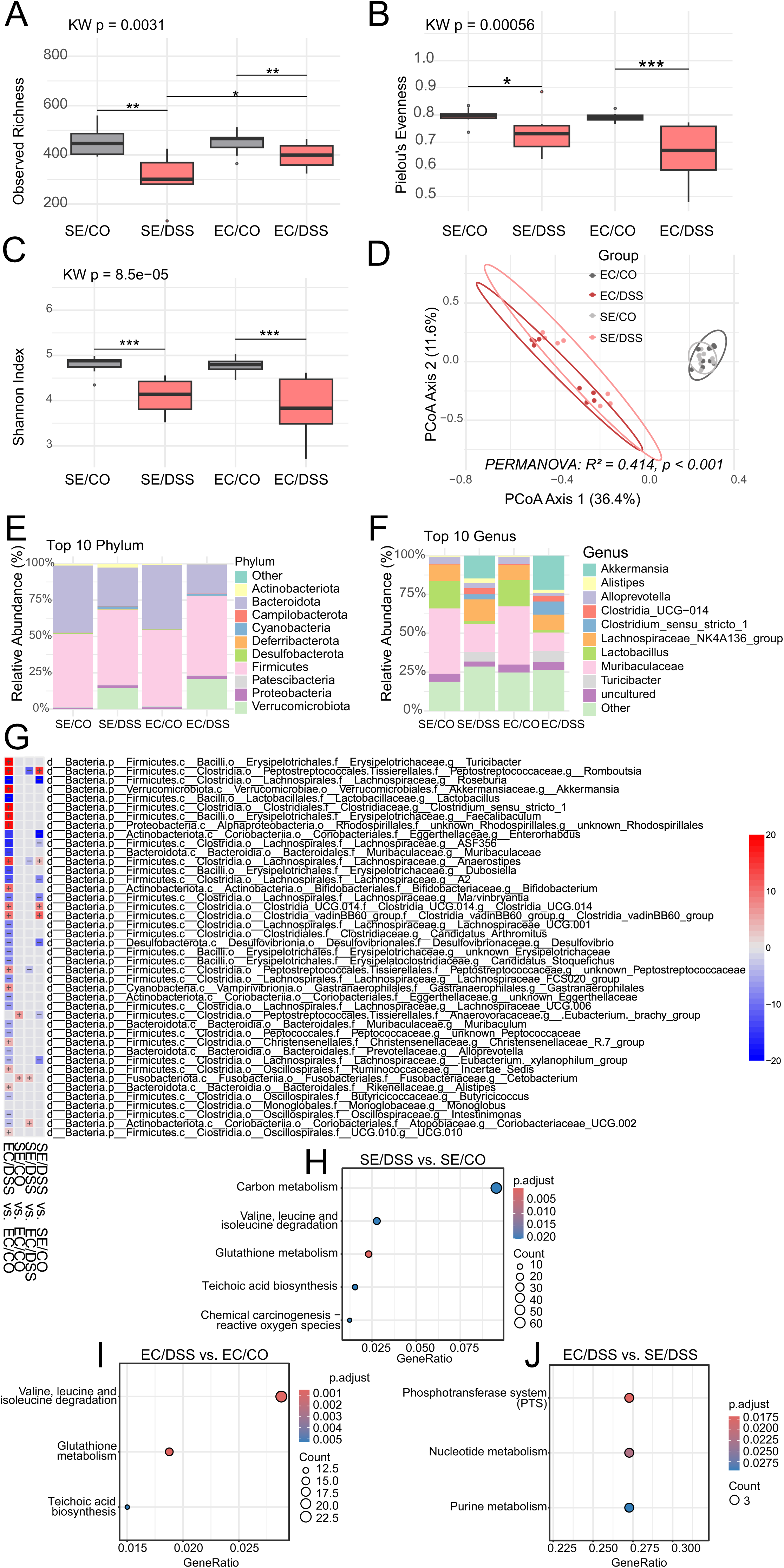
Microbial composition and colitis-induced shifts of the gut microbiota in standard environment and enhanced environmental complexity. Effects of enhanced environmental complexity (EC) and colitis on (A) alpha diversity: richness, (B) alpha diversity: Shannon index and (C) alpha diversity: evenness. Kruskal Wallis test followed by Wilcoxon rank-sum test. ***p<0.001, ** p<0.01, *p<0.05. n = 8-10/group. (D) PCoA plot displaying beta diversity based on Bray-Curtis dissimilarity. PERMANOVA. n = 8-10/group. (E) Taxonomic composition at the phylum level as shown by the relative abundance of the top 10 phyla; (F) Taxonomic composition at the genus level as shown by the relative abundance of the top 10 genera; (G) Heatmap of statistically significant different taxa between the experimental groups based on MaAsLin2 analysis; (H) Statistically significantly altered KEGG pathway terms between the SE/DSS and the SE/CO group; (I) Statistically significant altered KEGG pathway terms between the EC/DSS and EC/CO group; (J) Statistically significant altered KEGG pathway terms between the EC/DSS and SE/DSS group. CO = control; DSS = dextran sulphate sodium; EC = enhanced environmental complexity; SE = standard environment. n = 8-10/group.

Interestingly, some of the microbial alterations due to colitis were modified by EC. Most notably, the rise of *Akkermansia* and *Clostridium sensu stricto_1* and the reduction of *Alloprevotella* due to colitis appeared more pronounced in the EC/DSS group than the SE/DSS group (Fig. 8F). To test for significant differences in the abundance of all taxa at the genus level, we next performed differential expression analysis using MaAsLin2. In line with the observed difference in relative abundance of the most abundant genera, we found that colitis significantly alters the abundance of numerous taxa at the genus level (Fig. 8G). In detail, we found that *Romboutsia, Anaerostipses,* Clostridia UCG014 and Clostridia vadinBB60 group were significantly more abundant in both treated EC- and treated SE-housed mice compared to respective healthy EC- and healthy SE-housed mice, while *Eubacterium xylanophilum* group*, Desulfovibrio, Marvinbryantia,* Lachnospiraceae A2, Lachnospiraceae ASF356*, Enterorhabdus* and *Roseburia* were less abundant. Interestingly, however, many colitis-induced microbial alterations occurred only in EC and not in SE. Specifically, in the EC/DSS group we found a significantly higher abundance of *Turicibacter, Akkermansia, Clostridium sensu stricto 1, Faecalibaculum, unknown Rhodospirillales, Bifidobacterium,* unknown Peptostreptococcaceae, Gastranaerophilales, Christensenellaceae R.7 group, Ruminococcaceae Incertae Sedis, Rikenellaceae*, Alistipes* and Oscillospirales UCG.010 compared to the EC/CO group. In addition, *Lactobacillus, Muribaculaceae, Dubosiella,* Lachnospiraceae UCG.001*, Candidatus* Arthromitus, unknown Erysipelotrichaceae*, Candidatus* Stoquefichus, Lachnospiraceae FCS020 group, unknown Eggerthellaceae, Lachnospiraceae UCG.006*, Muribaculum,* unknown Peptococcaceae*, Alloprevotella, Butyricicoccus* and *Intestimonas* were less abundant in EC/DSS mice compared to EC/CO (Fig. 8G). In contrast, in SE-housed animals, a decrease in *[Eubacterium] brachy group* was the only SE-specific change due to colitis suggesting that EC facilitates colitis-induced microbial alterations. Importantly, several significant microbial shifts were also detected when directly comparing SE/DSS versus EC/DSS. This included a decrease in unknown Peptostreptococcaceae, *Anaerostipes*, and *Romboutsia* and an increase in Coriobacteriaceae UCG-002, suggesting the emergence of a distinct microbial signature in the EC/DSS group. Furthermore, *Cetobacterium* was less abundant in the EC/DSS group compared to SE/DSS, but also in healthy EC-housed animals relative to their SE-housed counterparts, indicating that this genus may represent a strain specifically affected by the EC housing condition. Taken together these data suggest that EC has an influence on colitis-induced microbial changes.

Functional prediction analysis using PICRUSt2 predicted marked potential differences in microbial metabolic across the experimental groups. In line with the differential abundance analysis these differences were most pronounced in the two colitis groups compared to their respective controls. Specifically, the comparison between SE/DSS and SE/CO revealed that 1226 KEGG Ortholog terms (KO) were predicted as significantly altered (Suppl. Table 11). KEGG pathway enrichment analysis revealed that carbon metabolism is the most represented pathway for this comparison (Fig. 8H). In addition to that, other significantly enriched pathways predicted are the degradation of amino acids such as valine, leucine and isoleucine, as well as glutathione metabolism, teichoic acid metabolism and reactive oxygen species. When comparing the EC/DSS group to EC/CO a total of 1525 KOs were found to be significantly different due to DSS treatment in the EC condition (Suppl. Table 12). Pathway analysis of these KOs revealed similar results like for the SE/DSS vs.SE/CO comparison including the terms valine, leucine and isoleucine degradation, glutathione metabolism and teichoic acid metabolism (Fig. 8I). In healthy animals, 29 KOs were predicted as significantly altered between the experimental groups, but no pathways emerged from the functional pathway enrichment analysis (Suppl. Table 13). When directly comparing EC/DSS with SE/DSS, we found that 19 KOs were significantly different between the groups (Suppl. Table 14). Pathway analysis revealed a predicted enrichment of phosphotransferase system (PTS), as well as purine and nucleotide metabolism (Fig. 8J).

### EC and colitis impair emotional-affective and social behaviour

Finally, in order to assess the neurobehavioural consequences of EC and DSS colitis, we performed a battery of behavioural tests to measure effects of the interventions on brain function (Fig. 9A). In the open field test (OFT), we found that colitis reduces the total distance travelled, independent of housing condition (DSS main effect: F_(1, 35)_ = 4.612; P=0.039; Fig. 9B). This effect was however more pronounced in EC/DSS mice, because EC itself also reduced locomotion independent of colitis (housing main effect: F_(1, 35)_ = 5.839; P=0.021; Fig.9B). EC also increased immobility in the OFT (housing main effect: F_(1, 35)_ = 6.562; P=0.015; Fig. 9C), with the highest average freezing times found again in the EC/DSS group, indicating heightened anxiety. We also assessed central zone time and the entries into the central zone, but did not observe significant differences between the experimental groups for these readouts (Fig. 9D-E). To investigate effects on anxiety behaviour in more detail, we also performed the elevated plus maze test. Here we did not observe significant differences in open arm entries between the treatment groups (Fig. 9F), but a significant interaction between DSS treatment and housing condition for the time spent on the open arms (DSS x housing interaction: F_(1, 35)_ = 7.456; P=0.010; Fig. 9G). Specifically, EC significantly reduced the time spent on the open arms in healthy mice, but this effect is absent in animals with colitis. In addition, DSS treatment reduced open arm time under standard conditions, but not in mice kept in EC. This suggests that both EC and colitis increase anxiety, but there is no synergistic effect of both interventions for this readout. To evaluate whether the interventions also influence social behaviour, we performed a social interaction test. Here we did not observe significant differences in the time spent in the social interaction (SI) zone (Fig. 9H), but animals treated with DSS entered the SI zone less frequently, indicating reduced sociality (main effect of DSS: F_(1, 35)_ = 4.976; P=0.032; Fig. 9I). Interestingly, the EC/DSS group showed the lowest number of entries, although this failed to reach statistical significance compared to the SE/DSS group.

**Figure 9.**
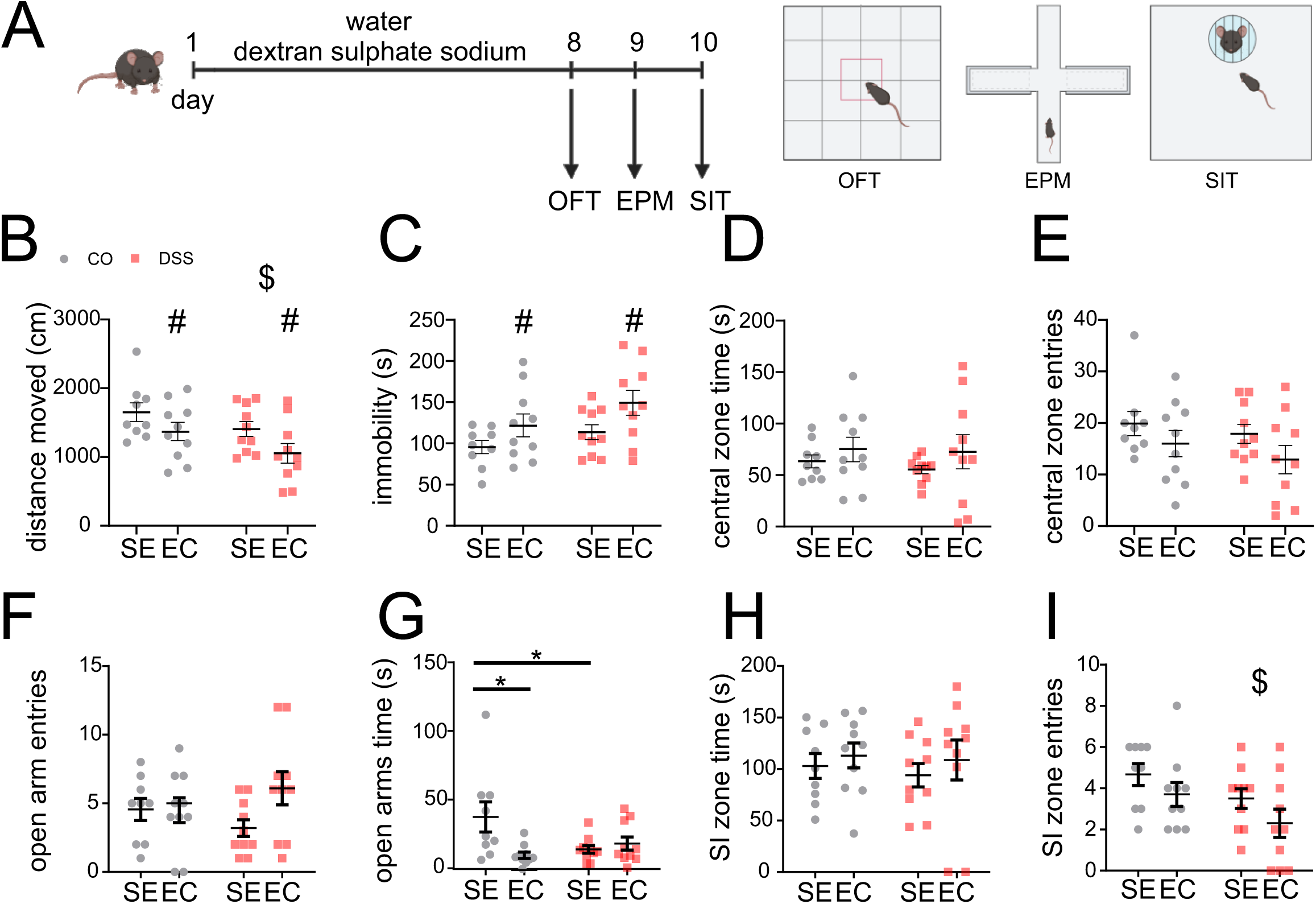
Behavioural effects of enhanced environmental complexity and colitis. A: Experimental timeline of dextran sulphate sodium (DSS) treatment (week 8 of differential housing) and schematic representation of behavioural tests performed during week 9. OFT: open field test; EPM: elevated plus maze test; SIT: social interaction test. (B) Total distance moved (cm) in the OFT; (C) Total time spent immobile (s) in the OFT; (D) Total time spent in the central zone (s) of the OFT; (E) Total number of entries in the central zone of the OFT; (F) Total number of entries into the open arms of the EPM; (G) Total time spent on the open arms (s) of the EPM; (H) Total time spent in the social interaction (SI) zone (s) of the SIT; (I) Total number of entries into the social interaction (SI) zone of the SIT. CO = control; DSS = dextran sulphate sodium; EC = enhanced environmental complexity; SE = standard environment. Data are presented as mean ± SEM. Two-way ANOVA followed by Tukey post-hoc test in case of a significant interaction. main effect of DSS: ^$^p < 0.05. main effect of EC: ^#^p < 0.05. Tukey post-hoc test: *p<0.05; n = 9-10/group.

## Discussion

IBD and other forms of gut inflammation are known to affect microbiota-gut-brain axis signalling, with detrimental consequences not only for the gut, but also for the brain (Bernstein *et al*. 2019; Bisgaard *et al*. 2022; Massironi *et al*. 2025; Petracco *et al*. 2025). Interestingly, however, this bidirectional communication between the gut and the brain can be modified by environmental factors such as chronic stress (Bisgaard *et al*. 2022; Bonaz *et al*. 2024). In the current manuscript our aim was to study the effects of EC, an environmental factor that is considered beneficial for laboratory mice (van Praag *et al*. 2000; Nithianantharajah and Hannan 2006), in a murine model of ulcerative colitis. For this, we characterized the effects of EC on all levels of the gut-brain axis including the gastrointestinal microbiota, gastrointestinal tissue, systemic circulation, brain tissue, brain interstitial fluid as well as cerebrospinal fluid. Against expectations, we found that EC had a sex-specific negative impact on experimental colitis. Female mice with colitis kept in EC had a more severe disease course and signs of enhanced peripheral inflammation compared to mice kept in SE. These differences were accompanied by metabolite alterations along the gut-brain axis, compartment-specific changes in immune cell compositions, behavioural alterations as well as changes to the gastrointestinal microbiome.

It is widely accepted that EC is a meaningful refinement procedure, often used as part of the 3Rs in animal research (Kulpa-Eddy *et al*. 2005). Especially in the field of animal behaviour, brain health and neuropsychiatric disease models, numerous studies have found a positive effect of this intervention (van Praag *et al*. 2000; Nithianantharajah and Hannan 2006). In addition, previous studies investigating the effects of EC on immunological processes in the brain have also detected various immune modulating effects. For example, EC has been found to promote the recruitment of T cells to the CNS and to activate microglial cells (Ziv *et al*. 2006). In disease models, EC can promote T-cell infiltration into the brain as well as faster virus clearance and disease resolution in a model of viral encephalitis (de Sousa *et al*. 2011) and EC can reduce neuroinflammation in experimental models of Alzheimer’s disease (Xu *et al*. 2016) However, much less is known about the effects of EC in the context of peripheral inflammation and how EC affects gut-brain axis signalling in this scenario. Previous studies investigating peripheral immunomodulating effects of EC have found that EC enhances splenic NK cell activity (Benaroya-Milshtein *et al*. 2004), increases the number of splenic B and T lymphocytes (Gurfein *et al*. 2014) and improves the chemotactic and phagocytotic functions of peritoneal leukocytes (Arranz *et al*. 2010), suggesting peripheral immune activating effects. In the current study, we found that EC leads to more pronounced colitis-induced tissue damage, enhanced weight loss and also a higher disease activity consistent with the mentioned immune activating effects of EC. Therefore, we hypothesised that the negative effects of EC are driven by alterations in immune cell populations along the gut-brain axis that have been previously linked to DSS-colitis and UC (Perše and Cerar 2012).

However, in the colon, the main site of DSS-induced inflammation, we did not observe any significant changes in the colitis-induced rise of various major immune cell populations by EC. This was also the case in the circulation and the brain, where colitis increased or decreased the number of multiple immune cell populations, with no additional change induced by EC. This suggests that changes in the number of the investigated immune cells was not the driving force for the disease exacerbating effects of EC in our model. However, it’s interesting to highlight that we found that EC lowered the number of neutrophils, T helper cells and activated microglia cells in the brain, independent of colitis, which is in line with previously described anti-inflammatory effects of EC in brain tissue (de Sousa Fernandes *et al*. 2022; Gonçalves *et al*. 2026). This might be a neuroprotective effect of EC through the strengthening of the blood-brain barrier (Paton et al., 2026). However, it might also be a consequence of sustained stress experienced by the animals kept in EC, as indicated by the high circulating corticosterone levels in the EC/DSS group. Given that stress and long-term exposure to corticosteroids can suppress immune system activity and basal choroid plexus gateway activity, limiting the recruitment of immune cells to the brain (Kertser *et al*. 2019), EC might have had negative effects for resolving CNS neuroinflammatory events due to colitis. Apart from these effects on immune cells in the brain, EC had much more pronounced effects on circulating and CNS metabolites. Among the several cytokines measured that have been previously linked to UC severity and activity (Nakase et al., 2022), we found that EC significantly lowered IL-22 levels, but increased IL-6 levels. This is of interest, because IL-22 has previously been shown to protect mice from experimental colitis (Zenewicz *et al*. 2008) and IL-6 is known as a major pro-inflammatory cytokine driving inflammation in IBD and also experimental colitis (Mudter and Neurath 2007). The high IL-6 levels and low IL-22 concentrations in the EC/DSS group thus support the view that EC can favour a pro-inflammatory environment during colitis.

Metabolome analysis of the gut-brain axis revealed numerous biological pathways affected by both EC and colitis. In line with previous studies suggesting that EC fosters a favourable metabolic milieu in healthy mice (Eduarda da Silva Fidélis *et al*. 2025), the EC/CO group showed compartment specific changes indicating reduced systemic inflammation and maintained CNS homeostasis. For example, plasma α linolenic acid (ALA; 18:3 n 3) was decreased in healthy EC mice, while it was elevated in brain and CSF, suggesting greater CNS availability of an anti inflammatory lipid precursor with neuroprotective effects due to EC (Yuan *et al*. 2022). In addition, tiglylcarnitine was increased in CSF and plasma of healthy mice kept in EC, but decreased in brain and ISF. Interestingly, lower levels of tiglylcarnitine in the brain have been associated with a protective genotype for Alzheimer’s disease (AD, Borkowski et al., 2025), while circulating levels of acylcarnitines were found to be reduced in AD and schizophrenic patients (Huo *et al*. 2019), supporting the view of protective EC effects via this pathway. In contrast, in EC mice exposed to DSS, the metabolome shifts toward a less favourable state. For example, uracil was reduced in brain, CSF, and plasma, suggesting dysregulated pyrimidine turnover with potential cytotoxic/pro inflammatory consequences (Löffler *et al*. 2005). Similarly, hypoxanthine was reduced by DSS treatment in brain, CSF, and plasma, consistent with altered purine metabolism and increased turnover (Tran *et al*. 2024). Moreover, tiglylcarnitine showed the opposite compartmentalization than in healthy EC mice – increased in brain and ISF but decreased in CSF.

A particularly noteworthy finding of our metabolomics analysis is that many microbiota-derived metabolites were altered between mice with colitis kept in SE or EC. Specifically, deoxycholic acid (DCA) and other secondary bile acids levels were elevated in EC/DSS mice. Because deoxycholic acid (DCA) and other bile acids can exert pro-inflammatory effects and have been associated with exacerbated Parkinson disease-like phenotypes in mice, their elevation in EC/DSS mice is consistent with phenotype worsening (Zhao *et al*. 2025). We also detected higher TMAO in the brains of EC/DSS animals, reflecting increased microbial trimethylamine (TMA) production and hepatic conversion to TMAO, indicating that EC facilitated TMA generation during experimental colitis. (Liu and Dai 2020). Interestingly, increased TMAO has been shown to promote brain aging and cognitive impairment in mice (Li et al., 2018). Some of these changes in microbial metabolites were also reflected in the gastrointestinal microbiota composition, pointing towards specific microbiota involved in the metabolite alterations observed. Specifically, we found that *Peptostreptococcaceae* were more abundant in the EC/DSS group than in the SE/DSS group, a family of bacteria that play a crucial role in the metabolism of primary bile acids into secondary bile acids (Larabi *et al*. 2023), likely contributing to the observed increase in DCA levels. In addition, some *Peptostreptococcaceae* and *Rombutsia* strains have been implicated in TMA production (Rath *et al*. 2017), which can explain the higher TMAO levels observed in the EC/DSS group.

Apart from these findings, the analysis of the microbial community revealed that several taxa shifted with DSS exposure independent of housing condition, but many additional taxa differed uniquely between EC/DSS versus its EC control. Among them, several genera typically associated with positive functions were reduced in the EC/DSS group (e.g., *Butyricicoccus*), whereas taxa linked to dysbiosis or disease (e.g., *Rhodospirillales*) were increased, supporting a state of EC specific dysbiosis under colitis (Devriese *et al*. 2017; Huangfu *et al*. 2021). Conversely, we also found evidence that some “beneficial” taxa were selectively enriched in EC/DSS mice (e.g., *Akkermansia*), and that others associated with adverse states like *Lachnospiraceae UCG 001* were less abundant (Khalili *et al*. 2024; Hwangbo *et al*. 2025). These bidirectional shifts of beneficial and detrimental bacterial taxa may suggest that EC alters niche availability and host factors such as mucus utilization and immune system activity. The EC/DSS microbiome profile shows a mixture of compensatory changes and opportunistic expansion of taxa adapted to the inflamed niche, emphasising the need for species/strain level resolution microbiome profiling in future studies, in order to better assess disease effects. Functional prediction using PICRUSt2 revealed that EC in the context of colitis was associated with enrichment of the phosphotransferase system, nucleotide metabolism, and purine metabolism, suggesting altered carbohydrate uptake/regulation and anabolic nucleotide turnover among taxa favoured in EC (Saier 2015), which can be tested directly in future studies using shotgun metagenomics, targeted metabolomics or further functional validation.

Given these characteristic microbial and metabolic shifts, as well as heightened disease activity in the EC/DSS group, we finally analysed how these changes affect animal behaviour. UC has been increasingly recognized as connected to a number of psychological comorbidities, reflecting the interaction between chronic gastrointestinal inflammation and mental health (Walker *et al*. 2008; Bernstein *et al*. 2018; Petracco *et al*. 2025). It has been previously found that DSS colitis is also linked to behavioural abnormalities in mice, such as enhanced anxiety, reduced social interaction, depressive-like symptoms and altered stress responses (Hassan *et al*. 2014; Reichmann *et al*. 2015; Komoto *et al*. 2022; Petracco *et al*. 2025). In line with previous studies, we observed that DSS colitis reduces total locomotion in the open field test, an expected sign of sickness behaviour (Hassan *et al*. 2014; Nyuyki *et al*. 2018), but this effect was more pronounced in EC/DSS animals reflecting the higher disease activity in this group. In addition, the EC/DSS animals were the most immobile in the open field test, suggesting heightened anxiety. This might be a consequence of higher stress hormone levels in the EC/DSS group, because elevated corticosterone levels are well known for their anxiety-inducing effects (McEwen 2007), but also of the described microbial and metabolic shifts. For example, the higher TMAO levels that we detected in the EC/DSS group, have been linked to anxiety behaviours and post-stress traumatic disorder (Romano *et al*. 2017; Baranyi *et al*. 2021), neuroinflammation and astrocyte activation (Brunt *et al*. 2020) and enhanced BBB permeability (Hu *et al*. 2024).

## Conclusion

In conclusion the current study suggests that EC, in the context of DSS colitis, is a chronic stressor, which exacerbates peripheral inflammation while limiting CNS immune cell recruitment in female mice. (Kalliokoski *et al*. 2010; Kertser *et al*. 2019)Moreover, EC in addition to DSS reshapes the gastrointestinal microbiome towards a less favourable state and alters metabolite dynamics across the gut-brain axis, favouring inflammation and anxiety-like behaviour. From a translational perspective, this study supports the view that environmental factors are key determinants of the disease course during chronic gastrointestinal inflammation such as IBD and IBD-associated neurobehavioral comorbidities.

## Materials and Methods

### Ethic statement

All experiments were approved by an ethical committee at the Federal Ministry of Science, Research and Economy of the Republic of Austria (permit number: GZ: 2021-0.878.133). All procedures were conducted according to the Directive of the European Parliament and of the Council of 22 September 2010 (2010/63/EU).

### Animals and study design

Before experiments, four-week-old male and female C57BL/6 mice (Charles River, Germany) were co-housed for 4 weeks in cohorts of 20 mice and habituated to the local animal facility of Medical University of Graz. The temperature of housing rooms was set at 21°C and humidity at 50%. They were maintained on a 12-hour light/dark cycle, with lights on at 6:00 AM and off at 6:00 PM and fed a standard mouse diet. After this habituation period, mice were randomly allocated to standard environment (SE) cages (length x width x height: 36.5×20.7×14.0 cm) or enhanced environmental complexity (EC) cages using Microsoft Excel’s RAND() function. Mice were kept in these differential housing conditions in groups of 5 mice per cage for 9 weeks, as previously described with minor modifications (Reichmann *et al*. 2013). Briefly, EC consisted of a large cage (length x width x height: 59.0×38.0×20.0 cm) containing enrichment objects that were regularly exchanged with each other (Suppl. Table 15) and mice had continuous access to a running wheel (Reichmann *et al*. 2013). For the whole period mice from both groups had access to water and food ad libitum. Experimental colitis was induced by administering 1.5% DSS (dextran sulphate sodium, MP Biomedicals, France; w/v in tap water) to female mice or 1% DSS to male mice via the drinking water for 7 days. Depending on the experiment, animals were euthanised either on day 9 or day 12 after the start of DSS treatment.

For the metabolomics experiments female mice were kept in EC or SE for 12 weeks. After week 10 of differential housing, mice underwent stereotaxic implantation of a cerebral flow microperfusion (cOFM) probe (cOFM-GD-2-2, JOANNEUM RESEARCH, Austria) into the striatum of the left hemisphere (0.6 / 2.2 / 4; AP/MD/DV) (Hummer *et al*. 2019; Altendorfer-Kroath *et al*. 2023). During the procedure, the animals were anaesthetised using inhalation anaesthesia (isoflurane at 1.5% in pure oxygen at 2 L/min, Abbott, Germany). Pain treatment during surgery was maintained by an opioid pain killer (fentanyl, 5µg/kg, i.p., Hameln Pharma, Germany). The cOFM probe was attached to the skull with UV-light curing dental acrylic and anchor screws. Postsurgical pain management involved presurgical injection of a NSAID (carprofen, 5 mg/kg, i.p., Zoetis, USA) and was maintained over three days after the surgery. To promote implant stability and optimal trauma healing, mice were then single-housed for 7 days with daily monitoring of general health, surgical site, and implant integrity (isolation period), followed by an additional 7 days in SE or EC housing without manipulation to allow complete restoration of blood–brain barrier (BBB) integrity (week 11). Dextran sodium sulphate (DSS) treatment started only after this 14-day postsurgical intervention (week 12) to minimize confounding from transient BBB permeability. Control animals were handled identically and received water only. ISF samples were collected on day 9 after the 14-day recovery period. For this, animals were slightly sedated (isoflurane, 0.8% in pure oxygen at 2 L/min, Abbott, Germany), not longer than 10 minutes, to be connected to a tethering system (Raturn, BASi, USA) allowing ISF sample collection from awake animals. ISF samples were collected hourly, over 24 hours at a flow rate of 1 µl/min. After the final ISF sample was collected, animals were disconnected from the cOFM system and deeply anesthetized with pentobarbital sodium (100 mg/kg, i.p.; Richter Pharma, Austria). CSF was collected from cisterna magna by puncturing the foramen magnum with a micropipette. Prior to tissue harvest, animals were transcardial perfused with PBS X1 + 10% heparin for approximately two minutes.

### Disease Activity Index Scoring

To assess the severity of experimental colitis a disease activity index was calculated, as previously described with minor modifications (Reichmann *et al*. 2015). The parameters considered were: weight loss (values: 0-5), presence of blood in the faeces or the perianal area (PAR) (values: 0-2) and stool consistency (values: 0-2) (Table 1).

**Table 1.** Disease activity index (DAI) scoring table used to assess the diseased phenotype of the experimental animals (score 0-9).

| parameter | score |  |
| --- | --- | --- |
| weight | 0 | gain > 1g |
|  | 1 | gain < 1 g |
|  | 2 | loss < 1 g |
|  | 3 | loss 1-3 g |
|  | 4 | loss 3-5 g |
|  | 5 | loss > 5 g |
| haemoccult test and perianal area (PAR) | 0 | neg. haemoccult test; no visible blood |
|  | 1 | pos. haemoccult test; no visible blood |
|  | 2 | visible blood in PAR |
| stool consistency | 0 | formed |
|  | 1 | soft |
|  | 2 | loose/no stool |

### Liquid consumption

Liquid intake over the DSS treatment period was measured by weighing drinking bottle weights every other day. For each cage, the total consumption during the DSS treatment period was calculated and divided by the number of animals per cage to obtain average liquid intake per animal. This was then normalized to each animal’s body weight.

### Hematoxylin and eosin (H&E) staining

Following euthanasia at day 12 after the start of DSS treatment, colons were harvested, flushed with cold PBS, and cut longitudinally. The medial portion of the colon was processed into swiss rolls and fixed in 4% neutral-buffered formalin for 48-72 hours at 4°C. Tissues were then dehydrated, embedded in paraffin, and sectioned at 5 μm thickness using a microtome. Sections were mounted on glass slides (Epredia, USA) and stained with hematoxylin and eosin (H&E) using standard protocols. Briefly, slides were deparaffinized in xylene, rehydrated through graded ethanol, stained with hematoxylin, rinsed in tap water, then counterstained with eosin. After dehydration and clearing, slides were coverslipped with mounting medium (Roti®Histokitt II, Carl Roth, Germany). Images of H&E-stained sections were acquired using a light microscope (Olympus BX53F2, Olympus, Japan), and histological scoring of 6 images per animal was performed in a blinded manner to assess crypt architecture, goblet cell depletion, muscle thickening and immune cell infiltration according to established criteria (scoring 0-10) adapted from Kim et al., 2012.

### MPO ELISA

Colonic myeloperoxidase (MPO) content was measured in the distal portion of snap frozen colons collected at day 9 after the start of DSS treatment, as previously described (Reichmann et al. 2013). Briefly, 10 mg from each sample was taken and homogenized with a Precellys Evolution Touch Homogenizer (Bertin Technologies, France) in 250 µl lysis buffer (200mM NaCl, 5mM EDTA, 10mM Tris Pufferan®, 10% Glycerin dissolved in distilled water) supplemented with Ready Shield Protease Inhibitor Cocktail (Sigma-Aldrich, Germany). The homogenates were diluted 1:10 and MPO ELISA was performed according to manufacturer’s instruction (Hycult Biotech, USA).

### Flow cytometry

For colonic tissue, single cell suspensions of the whole colon were prepared to isolate intra-epithelial leukocytes (Kienzl *et al*. 2020). The colons were cut in small pieces and digested with disodium EDTA (2 mM; Merck Group, Germany) and DL-dithiothreitol (0.15 mg/mL; Sigma-Aldrich, Germany) in HBSS (Hanks’ Balanced Salt Solution; Thermo Fisher Scientific, USA) for 40 minutes at 37°C while rotating at 250 rpm. Then, the tissue was flushed through a 100 µm strainer and washed with PBS (phosphate buffered saline; Thermo Fisher Scientific, USA). Blood samples were processed first by lysing red blood cells with RBC Lysis Buffer (9000 mg/mL Ammonium Chloride, 1000 mg/mL KHCO3, 0.37 mg/mL EDTA in distilled water) for 5 minutes on ice. Then they were washed with PBS. After washing, cells were counted and used for antibody staining (Table 2). Brains were collected from the animals after perfusion of their heart with PBS. To prepare single cell suspensions, the brains were manually minced and flushed through 100 µm strainers, then incubated with RBC lysis buffer and washed with PBS. After that, the tissue was centrifuged with Percoll® (Cytiva, USA) at room temperature and the myelin layer was discarded. The samples were flushed through 40 µm strainers strainer and washed with PBS. Cells were then counted, stained (Table 2), fixed with IC Fixation Buffer (eBioscience, USA) and acquired using a BD LSRFortessa flow cytometer, with data collection performed in FACSDiva (BD Biosciences, USA). Data analysis and compensation were carried out using FlowJo 10 (BD Biosciences, USA). Fluorescence minus one (FMO) controls were used to establish gating boundaries. Gating strategies are shown in Supplementary Figures 3, 4.

**Table 2.** Antibodies used for staining the single-cell suspensions obtained from colon, blood and brain samples.

| Antibody | Company | RRID |
| --- | --- | --- |
| CD45-BV785 | BioLegend | AB_2564590<br>(BioLegend<br>Cat. No.<br>103149) |
| CD3-PECy7 | Thermo Fisher | AB_469572 |
| CD4-BUV496 | BD Biosciences | AB_2813886 |
| CD8-PerCPCy5.5 | BioLegend | AB_2075238<br>(BioLegend<br>Cat. No.<br>100734) |
| NKp46-BV510 | BioLegend | AB_2563290<br>(BioLegend<br>Cat. No.<br>137623) |
| CD11b-BUV737 | BD Biosciences | AB_2738811 |
| Ly6C-FITC | BioLegend | AB_1186134<br>(BioLegend<br>Cat. No.<br>128005) |
| Ly6G-PE/Dazzle594 | BioLegend | AB_2566319<br>(BioLegend<br>Cat. No.<br>127648) |
| CD11c-BV605 | BioLegend | AB_2562415<br>(BioLegend<br>Cat. No.<br>117334) |
| F4/80-BUV395 | BD Biosciences | AB_2739304 |
| SiglecF-PE | BD Biosciences | AB_394341 |

### Corticosterone ELISA

Plasma corticosterone concentrations were measured using the Corticosterone ELISA Kit (Enzo Life Sciences, Cat. No. ADI-900-097) according to the manufacturer’s instructions as previously described (Reichmann et al. 2015). Briefly, plasma samples were diluted 1:200 with the provided assay buffer and loaded in duplicate into the 96-well plate following the assay protocol. Absorbance was read at 405 nm using a microplate reader and corticosterone concentrations were calculated from the standard curve using a 4-parameter logistic (4PL) curve-fitting model.

### Procartaplex Multiplex ELISA Assay

A custom magnetic bead-based immunoassay was used to quantify cytokine levels in plasma samples collected at day 9, employing the ProcartaPlex multiplex suspension array system (ThermoFisher Scientific, USA) based on Luminex xMAP technology. The assay was performed according to the manufacturer’s protocol. Briefly, plasma samples were thawed on ice, centrifuged at 10,000 × g for 10 minutes at 4°C to remove debris, and diluted appropriately in assay buffer. Samples, standards, and quality controls were incubated with magnetic beads conjugated to capture antibodies specific to the selected cytokines. After washing, biotinylated detection antibodies were added, followed by streptavidin-phycoerythrin (PE). Plates were processed using a magnetic plate washer between each incubation step to remove unbound components. The beads were then analysed using the Bio-Plex 200 system, and data were acquired and processed using Bio-Plex Manager Software version 5.0. Cytokine concentrations were determined using a five-parameter logistic (5-PL) standard curve and expressed as fluorescence intensity (FI) (background-corrected). All samples were run in duplicates. The custom panel included the following cytokines: GRO-α, IL-18, IL-22, IL-6, TNF-α.

### Metabolomics measurements and analyses

Metabolome analyses were conducted on plasma, cerebrospinal fluid (CSF), interstitial fluid (ISF), and brain tissue samples. For the measurement of ISF, samples collected between 1 and 3 hours after the start of the OFM procedure were pooled and processed. Brain tissue samples (one brain half) were homogenized using a bead mill and extracted with cold methanol as described previously (Paulus *et al*. 2022), whereas biological fluids underwent overnight extraction in cold methanol as described previously (Monedeiro *et al*. 2025). In short, specified sample amounts were added to ice-cold methanol, vortexed, incubated for at least 8 h at -80°C, followed by a centrifugation step (10min, 14000g, 4°C). Supernatant was transferred to fresh sample tubes, evaporated until dry using nitrogen, and residue was resolved in acetonitrile (ACN)/water (50:50, v:v).

Metabolite profiling was performed using a Vanquish UHPLC system coupled to a Q Exactive high-resolution mass spectrometer (HRMS; Thermo Fisher Scientific, USA) operated in both positive and negative electrospray ionization modes, covering a mass range of 70–920 m/z. Chromatographic separation was achieved with an Acquity UPLC BEH Amide column (1.7 µm, 100 mm × 2.1 mm; 130 Å; Waters, USA) with an injection volume of 2 µL. Analyses were carried out using ammonium acetate (10 mM)/ACN (90:10, v:v) at pH 9 (adjusted with ammonium hydroxide) as the mobile phase A, and ammonium acetate (10 mM)/ACN (10:90, v:v) as the mobile phase B in a gradient elution as described in (Monedeiro *et al*. 2025). HRMS electrospray ionization parameters were as follows for both negative and positive mode: spray voltage 3500 V, capillary temperature 300 °C, sheath gas 18 a.u., auxiliary gas 9 a.u., sweep gas 0 a.u., s-lens RF level 65, with scan parameters being: resolution = 70,000, microscans = 1, automatic gain control target = 3 × 10^6^, maximum injection time = 250 ms. Each sample matrix was analysed in a single analytical run, with dedicated pooled quality control samples (QC) included per matrix to ensure reproducibility and data quality. QCs, blank samples (mobile phase B) and Ultimate Mix (UM, prepared as described in (Vogel *et al*. 2019)) were injected sequentially. To avoid time-dependent bias, sample analysis order was randomized in a stratified way, while analytical runs were subdivided into smaller sub-batches to limit autosampler residence time and reduce potential time-dependent sample degradation. Acetonitrile (≥99.9 %, AppliChem, Darmstadt, Germany) and methanol (≥99.8 %, VWR, Austria) were purchased as reagent-grade chemicals, certified for high-performance liquid chromatography (HPLC) use. All aqueous solutions were prepared using purified water (18.2 MΩ cm, Milli-Q; Merck, Germany).

A targeted data analysis was conducted using Skyline software (v. 23.1.0.455, MacCoss Lab, University of Washington, USA). Data curation, analysis, and visualization were performed in R (v4.2.1) using RStudio (v2023.06.2; Posit PBC, USA) and TIBCO Spotfire (v7.5.0). Metabolomics data quality was assessed using predefined thresholds for mass accuracy, retention time variability, signal variability and missing values, together with acceptable peak shape, minimal signal drift over measurement time or batch, low blank contribution in QCs, and absence of analytical outliers (Vogel *et al*. 2019; Monedeiro *et al*. 2025). Metabolomics data were either median-normalized and log10-transformed (brain tissue) or only log10-transformed (plasma, ISF and CSF). Normality was assessed primarily using Shapiro–Wilk test. Principal component analysis (PCA) was performed on centered and scaled data using R stats package; missing values were solely imputed for PCA using regularized expectation-maximization implemented in missMDA. Group-wise pairwise comparisons were conducted using linear models implemented using the nlme package. Pathway enrichment analysis was performed with FELLA using KEGG IDs of significantly altered metabolites (p < 0.05) for each contrast of interest. Enrichment was based on hypergeometric testing with false discovery rate correction and using *Mus musculus* KEGG pathway database as reference. Enrichment plots were generated using ggplot2.

### 16s rRNA Analysis

Microbial DNA was extracted from faecal samples obtained at the height of inflammation post-DSS treatment (day 8) using the PureLink™ Microbiome DNA Purification Kit (Invitrogen, USA) according to the manufacturer’s protocol. The V3–V4 region of the bacterial 16S rRNA gene was amplified using the following primers FW: CCTACGGGNGGCWGCAG and RV: GACTACHVGGGTATCTAATCC, and sequencing was performed on the Illumina MiSeq platform. Raw sequences were processed using QIIME2 (version 2023.2), including quality filtering, denoising, and chimera removal with the DADA2 plugin. Amplicon sequence variants (ASVs) were aligned and taxonomically classified using a pretrained Naïve Bayes classifier based on the SILVA v138 database. Subsequent analyses, including alpha and beta diversity, as well as taxonomic composition, were carried out and visualized using R (packages: phyloseq, vegan, ggplot, decontam, SRS, biomformat, dplyr, tidyr, ALDEx2, reshape2, tibble, rstatix). Differential abundance analysis across taxonomic levels (phylum, family, and genus) was conducted using the R package Maaslin2 with default settings and adjusting for "Cohort", "Extraction" and "Cage" (Mallick *et al*. 2021). PICRUSt2 analysis was performed using NAMCO (https://exbio.wzw.tum.de/namco/) and statistical analysis and visualisation were conducted with R (packages: LinDA (Zhou *et al*. 2022), ggplot2, dplyr, tibble, tidyr). Functional enrichment analysis was carried out using R/shiny package MicrobiomeProfiler.

### Open Field Test

The open field test (OFT) was performed using a 50 × 50 × 50 cm (length × width × height) opaque grey plastic box. Its area was divided into an inner zone measuring 36 × 36 cm and a surrounding outer zone, as previously described (Reichmann et al. 2016). Each mouse was placed in the middle of the box and behaviour was recorded for 5 minutes using a video camera positioned above its centre. The recordings were captured and analysed using Ethovision XT 17 software (Noldus, The Netherlands). The parameters measured were: time spent in the inner zone, number of entries into the inner zone, time spent in the outer zone, total distance covered in the OFT and total time spent immobile. An observed reduction in the inner zone time and entries was interpreted as an increase in anxiety-like behaviour. The OFT box was cleaned with water then 70% Ethanol after each test session.

### Elevated Plus Maze

The elevated plus maze test (EPM) was performed using a cross-shaped maze with two open arms (30 x 5 x 0.5 cm) perpendicular to two closed arms (30 x 5 x 0.5 cm), with a centre platform (5 x 5 x 0.5 cm), as previously described (Reichmann et al. 2016). Each mouse was placed at the intersection of the four arms and behaviour was recorded for 5 minutes using a video camera positioned above the maze. The recordings were captured and analysed using Ethovision XT 17 software (Noldus, The Netherlands). The parameters measured were: time spent in the open arms, number of entries into the open arms, time spent in the closed arms, total distance covered in the EPM and total time spent immobile. An observed reduction in the open arms time and entries was interpreted as an increase in anxiety-like behaviour. The EPM maze was cleaned with water then 70% Ethanol after each test session.

### Social interaction test

The social interaction (SI) test was performed as previously described (Hassan *et al*. 2014). A cylindrical meshwork container (7 × 10 cm, diameter × height) was positioned adjacent to one of the walls of the open field box and midway along its length. The area around it, 8 cm diameter, was defined as “social interaction zone”. The test mouse was placed on the other side of the box, near the wall opposite the cylindrical container, and was allowed to explore for 3 minutes before being returned to its home cage. Subsequently, an unfamiliar mouse (stimulus mouse) was introduced into the cylindrical container, and the test mouse was given an additional 3 minutes to explore the box. The recordings were captured and analysed using Ethovision XT 17 Software (Noldus, The Netherlands). The parameters measured were: time spent and number of entries in the social interaction zone, total distance covered in both sessions and total time spent immobile. The box was cleaned with water then 70% Ethanol after each test session.

### Statistical Analysis

Statistical analyses were conducted using GraphPad Prism 10 (GraphPad Software, USA). For comparisons involving one variable and two factors (housing condition and colitis), data were analysed by two-way analysis of variance (two-way ANOVA) evaluating main effects and interactions between the two factors. In case of a significant interaction term, post-hoc testing using Tukey’s post hoc test for multiple pairwise comparisons was performed (SE/DSS vs. SE/CO, EC/DSS vs. EC/CO, EC/CO vs. SE/CO, EC/DSS vs. SE/DSS). In case of a non-significant interaction term the main effects were interpreted and planned comparison of EC/DSS vs. SE/DSS was performed. Body weight changes over time were assessed using repeated-measures ANOVA, with time as the within-subject factor and treatment group as the between-subject factor. Kruskal–Wallis tests were used to compare microbiome-derived metrics. When the overall test was significant, pairwise post hoc comparisons were performed with Wilcoxon test appropriate correction for multiple testing (Benjamini-Hochberg). Unless otherwise specified, results are reported as mean ± SEM, and statistical significance was defined as P<0.05.

## Acknowledgments

We are grateful to Sabine Donner for technical assistance and would like to thank members of the biomedical research unit of the Medical University of Graz (BMF) for mouse care and maintenance. We also thank the Core Facility Molecular Biology (CF-MB) at the Medical University of Graz for their support regarding the 16s amplicon sequencing and Bioplexx measurements. Figures 1A and 9A were created in BioRender. Reichmann, F. (2026) https://BioRender.com/9d8rw5i and https://BioRender.com/dgrg0zk. This research was funded in whole or in part by the Austrian Science Fund (FWF) [10.55776/P35774]. For open access purposes, the author has applied a CC BY public copyright license to any author accepted manuscript version arising from this submission. GP and IF are funded by the FWF within the PhD program Molecular Medicine of the Medical University of Graz. Work in the lab of RS is funded by the Austrian Science Fund (FWF; PAT4791123)

## Declaration of generative AI and AI-assisted technologies in the manuscript preparation process

During the preparation of this work the authors used medunigraz.academic-ai.at in order to improve writing and for language editing, as well as chatgpt.com for coding tasks. After using these tools, the authors reviewed and edited the content as needed and take full responsibility for the content of the published article.

