## Supplementary figures and images for "Enhanced environmental complexity worsens experimental colitis and dysregulates microbiota-gut-brain axis signalling in female mice"

### Suppl. Figure 1

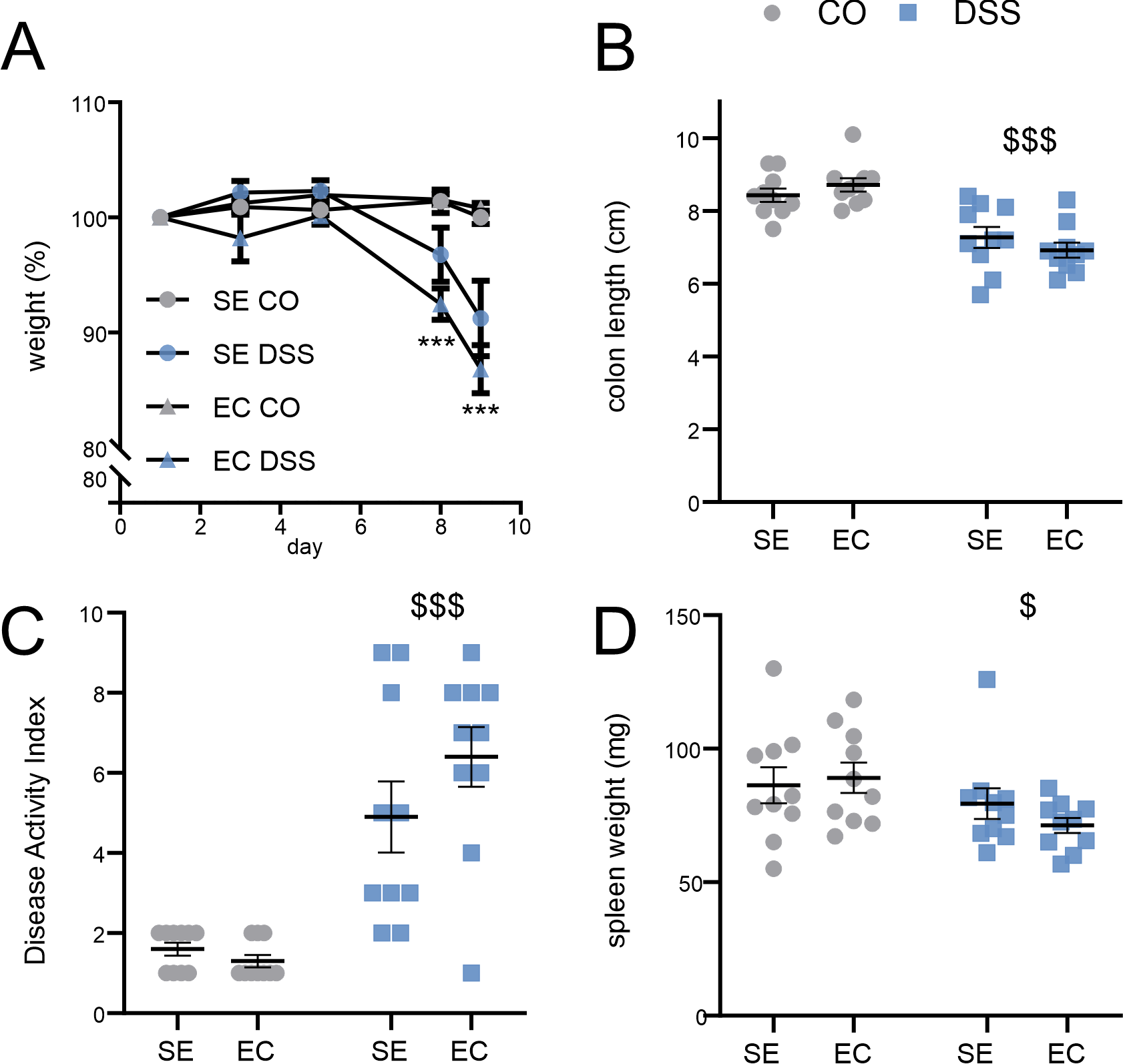

### Suppl. Figure 2

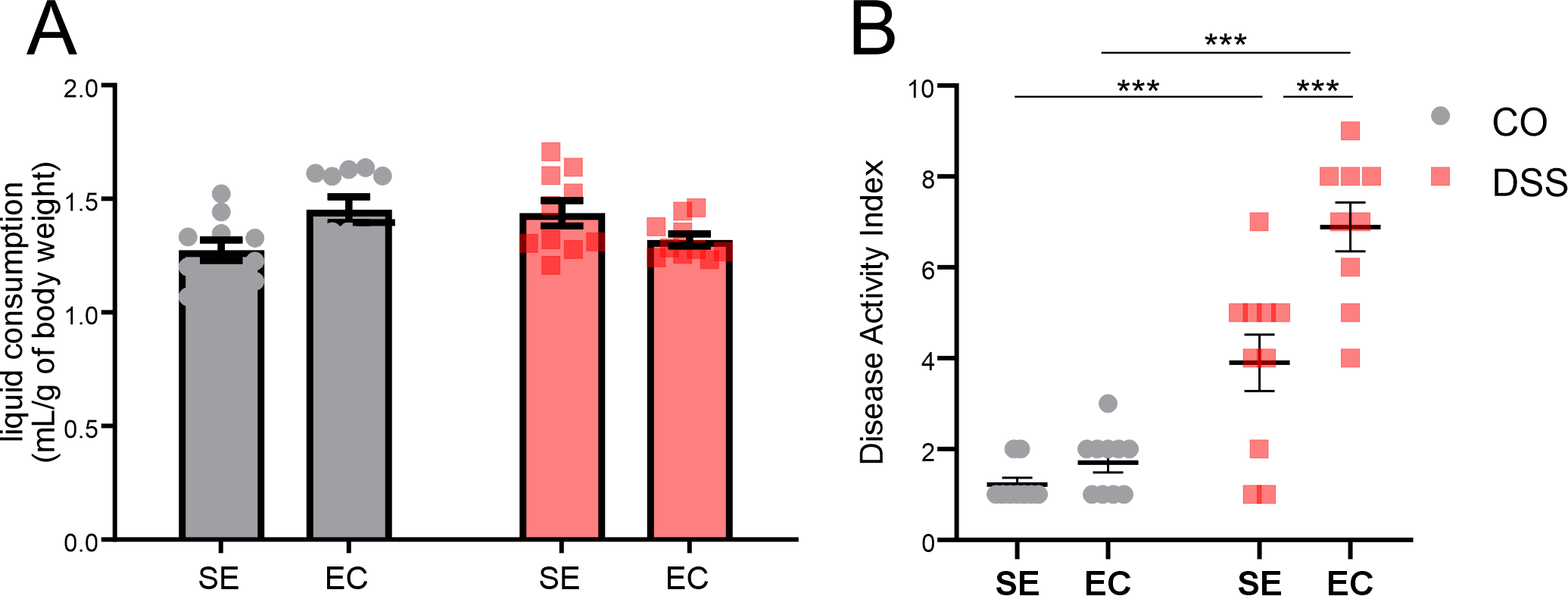

### Suppl. Figure 3

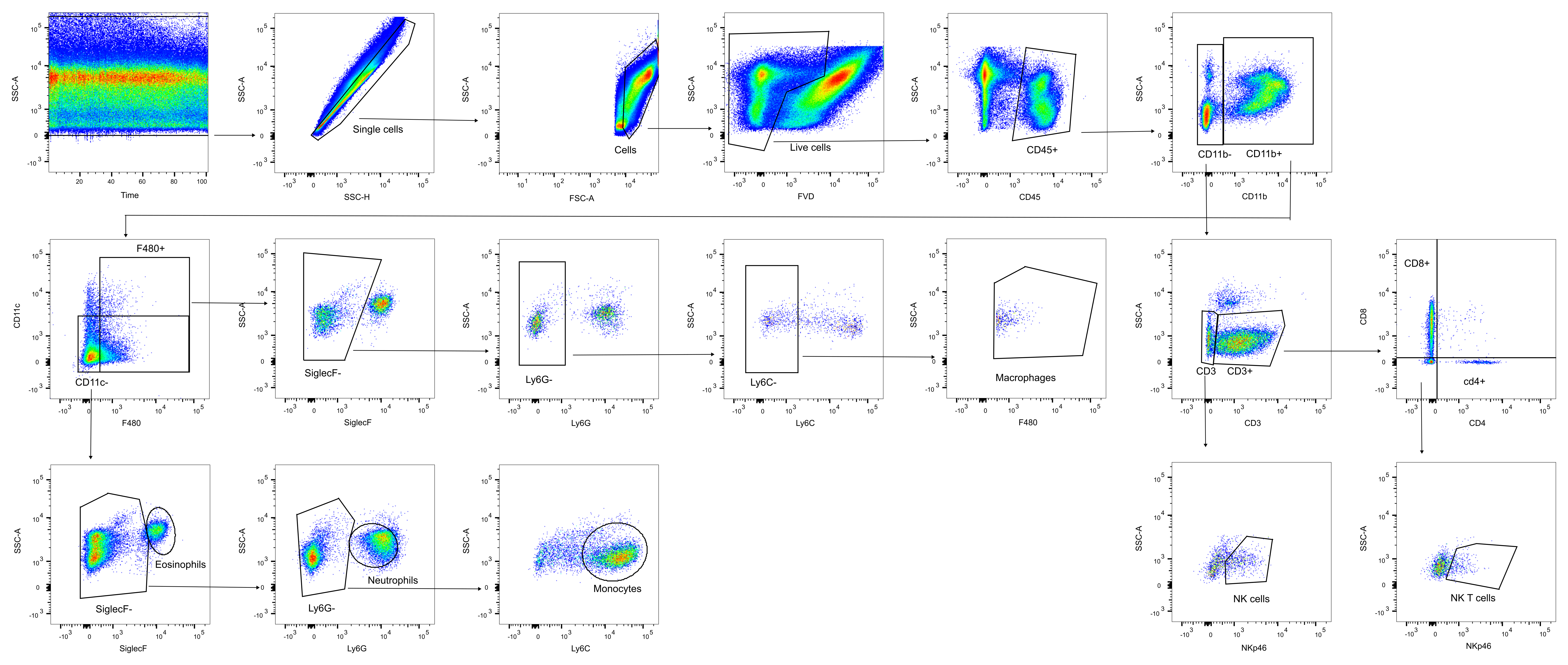

### Suppl. Figure 4

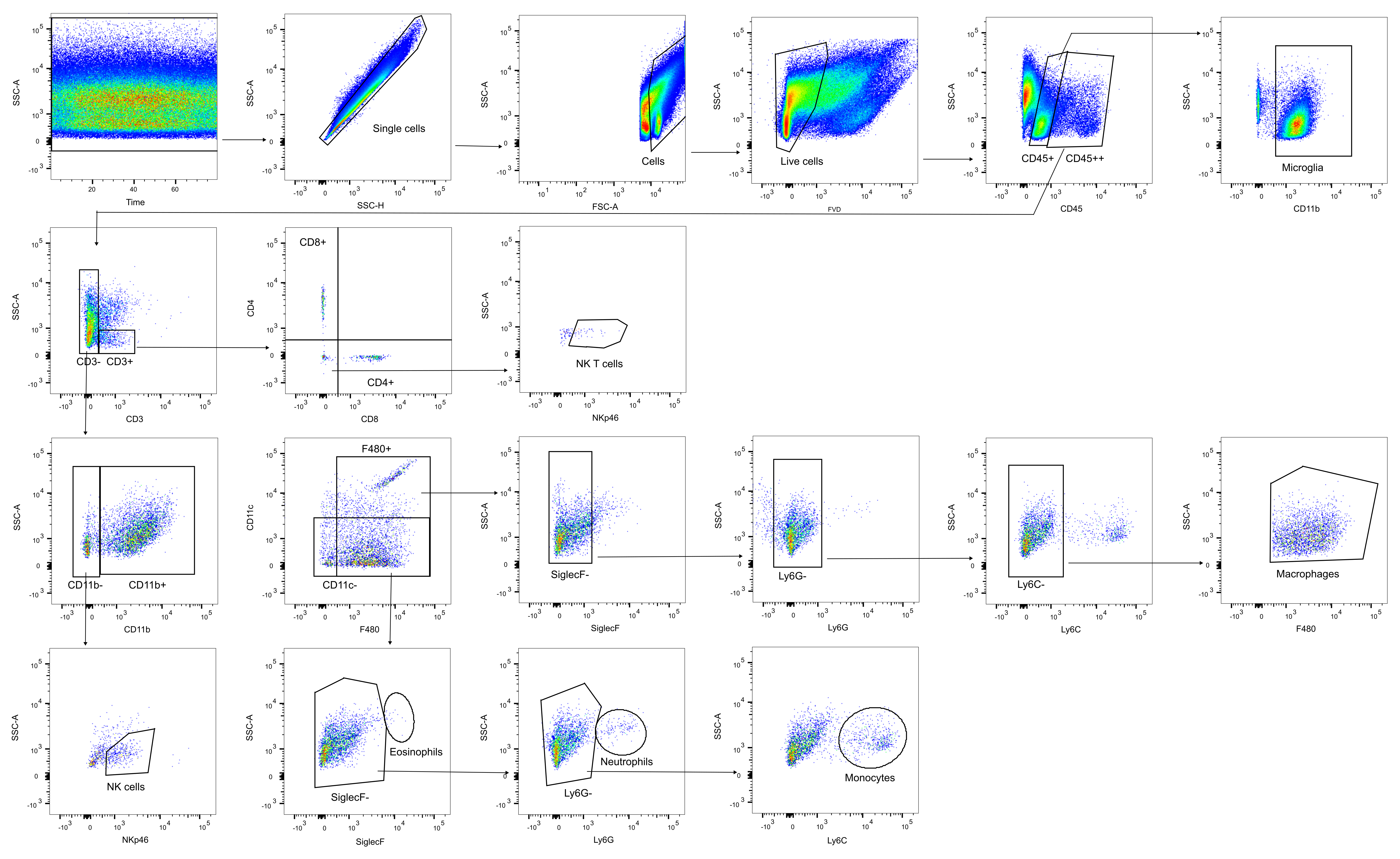

### Suppl. Figure 5

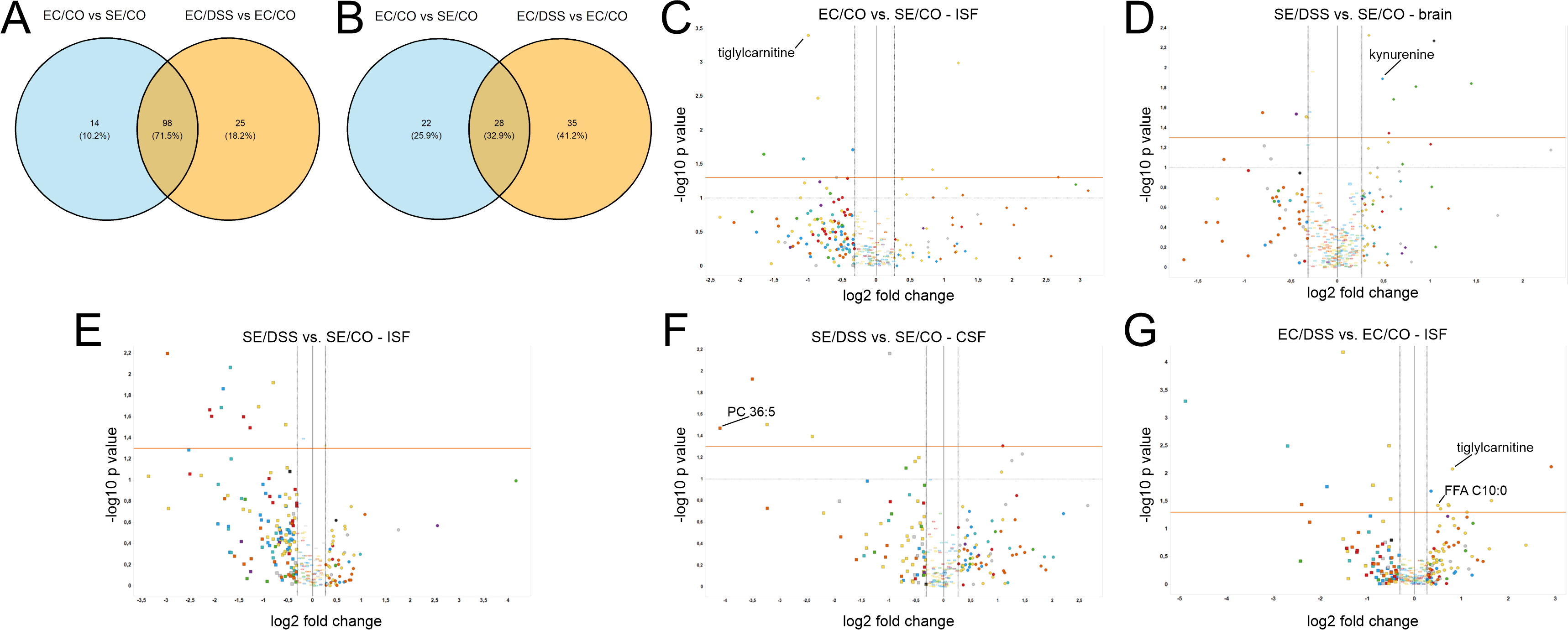
